# Opto-activation of cortical somatostatin interneurons alleviates parkinsonian symptoms

**DOI:** 10.1101/460535

**Authors:** Marie Vandecasteele, Sébastien Valverde, Charlotte Piette, Giuseppe Gangarossa, Willy Derousseaux, Asier Aristieta Arbelaiz, Jonathan Touboul, Bertrand Degos, Laurent Venance

## Abstract

Deep brain stimulation of the subthalamic nucleus is a symptomatic treatment of Parkinson’s disease but benefits only to a minority of patients due to stringent eligibility criteria. To investigate new targets for less invasive therapies, we aimed at elucidating key mechanisms supporting deep brain stimulation efficiency. Here, using *in vivo* electrophysiology, optogenetics and modeling, we found that subthalamic stimulation normalizes pathological hyperactivity of motor cortex pyramidal cells, while concurrently activating somatostatin and inhibiting parvalbumin interneurons. *In vivo* opto-activation of cortical somatostatin interneurons alleviates motor symptoms in a parkinsonian mouse model. A mathematical model highlights how the decrease in pyramidal neurons activity can restore information processing capabilities. Overall, these results demonstrate that activation of cortical somatostatin interneurons may constitute a less invasive alternative than subthalamic stimulation.

**One-sentence Summary:** Deep brain stimulation recruits cortical somatostatin interneurons, and their opto-activation is beneficial in Parkinson’s.

---

Parkinson’s disease results from the neurodegeneration of the nigro-striatal dopaminergic neurons. The main symptomatic treatment for Parkinson’s disease consists in substituting lacking dopamine with levodopa and dopaminergic agonists (1). However, after a typical “honeymoon” period with dopaminergic therapy, patients inevitably develop levodopa-induced motor complications. At this stage, deep brain stimulation at high frequency of the subthalamic nucleus (STN-DBS) constitutes to date the most efficient symptomatic treatment (2, 3). However, due to its surgical invasiveness and strict eligibility criteria, STN-DBS benefits only to a minority of patients (~5-10%). Several hypotheses have been proposed to explain the beneficial effects of STN-DBS *(4-6),* but the exact mechanisms remain elusive. Notably, a growing body of evidence points towards a cortical effect of STN-DBS in parkinsonian rodent models (7-11) and patients (12-17subpopulation across cortical). Here, we reasoned that mimicking the cortical effects of STN-DBS could reproduce its therapeutic benefits, thus paving the way for less invasive approaches. To do so, we i) determined STN-DBS effects on cortical cell-type specific populations using a combination of *in vivo* electrophysiological and optogenetic approaches, ii) reproduced these effects using optogenetics in freely-moving parkinsonian mice, and *iii*) explored mathematically how DBS and DBS-guided optogenetics could restore cortical information processing capabilities.

To understand how DBS affects cortical activity, we first performed *in vivo* single unit extracellular recordings of layer V motor cortex neurons in a rat model of Parkinson’s disease, the 6-hydroxydopamine (6-OHDA) lesion of the substantia nigra *pars compacta* (Fig. 1A). We observed an elevated spontaneous firing rate of primary motor cortex (M1) pyramidal cells in 6-OHDA-lesioned rats compared to sham animals (p=0.047) (Fig. 1B), in line with previous studies *(70, 78).* STN-DBS decreased spontaneous activity of pyramidal cells both in 6-OHDA-lesioned (p=0.021) and in sham (p=0.018) rats (Fig. 1B). The percentage of cells inhibited by STN-DBS (>70%) and the magnitude of the effect (two-fold decrease in firing rate) were similar (p=0.53, p=0.75) in 6-OHDA-lesioned and sham conditions (Fig. 1C). We next explored the mechanistic underpinnings related to the decreased activity of pyramidal cells under STN-DBS. For this purpose, we performed single-cell *in vivo* intracellular recordings of electrophysiologically identified M1 pyramidal cells in anesthetized rats (Fig. 1A and D). The decreased firing rate (p<0.001) of M1 neurons during STN-DBS was accompanied by a hyperpolarization of their membrane potential (p<0.001) (Fig. 1D). In cells that were antidromically activated by STN stimulation, we observed that the evoked spikes were rapidly shunted (Fig. S1A). We next investigated the effect of DBS on evoked activity by applying successive depolarizing current steps, and observed that DBS reduced the evoked firing rate of M1 pyramidal cells (Fig. 1E). This modification of the input-output gain function caused an increase of the rheobase (p=0.031), without affecting the gain (p=0.17) (Fig. 1E). To better characterize the evoked conductances in M1 pyramidal cells, we applied single stimulations in the STN (Fig. 1F and G). We observed a post-stimulation hyperpolarization, sufficient to delay the firing activity (p=0.0057) (Fig. 1F). Analysis of the voltage-dependency showed that the evoked postsynaptic response reversed around -71mV (Fig. 1G), which corresponds to the chloride reversal potential. These results suggest that STN-DBS recruits GABAergic circuits responsible for the hyperpolarization of pyramidal cells and the increase in rheobase, leading to the reduced activity detected in both extra-and intracellular recordings of M1 pyramidal cells. The short (<5ms) and low variability (SD<1ms) latency of the STN-evoked response (Fig. S1B) indicates a monosynaptic connection. The absence of direct inhibitory afferents from the STN to the cortex argues for a local recruitment of cortical GABAergic networks by an antidromically-evoked axonal reflex of cortico-STN pyramidal cells.

**Figure.1.**
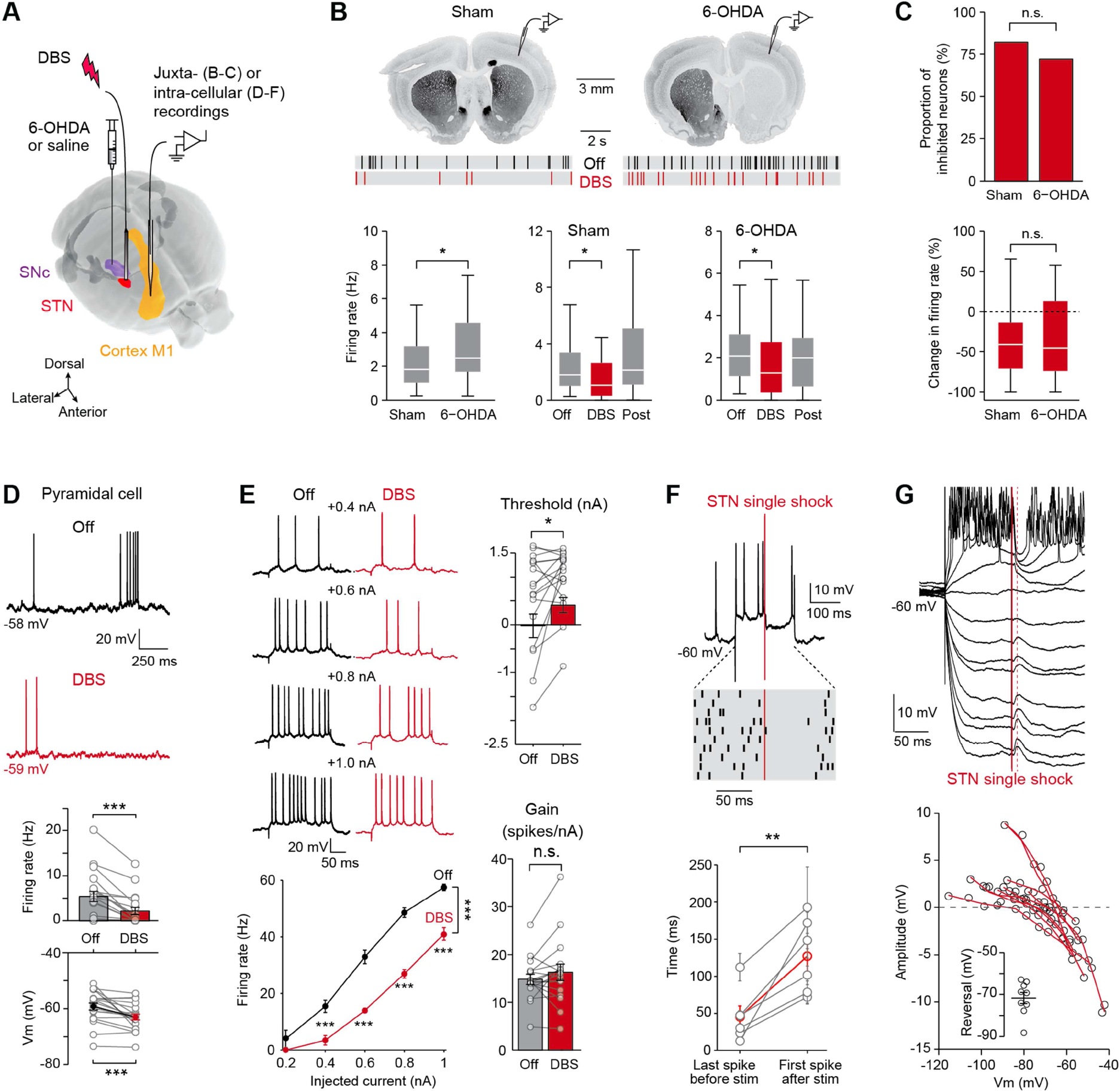
STN-DBS inhibits Ml pyramidal cells. (A) Experimental design. (B) TH immunostaining and raster plot of M1 neurons recorded in sham and 6-OHDA-lesioned rats. Spontaneous activity of cortical neurons increased in 6-OHDA-lesioned (n=46 neurons) compared to sham (n=43) rats (Mann-Whitney U test, p=0.047); STN-DBS decreased neuronal activity, in both sham-(Wilcoxon signed rank test, n=33, p=0.018) and 6-OHDA-lesioned (n=25, p=0.021) rats. (C) The proportion of inhibited neurons (Fisher’s exact test, p=0.53) and the magnitude of the effect (Mann-Whitney U test, p=0.75) are similar in sham-and 6-OHDA-lesioned rats. (D) M1 pyramidal neurons recorded intracellularly display a decreased activity (paired t-test, *n=20,* p<0.001), and hyperpolarized membrane potential (paired t-test, p<0.001) during STN-DBS. (E) Spiking activity evoked in M1 pyramidal neurons. *f-I* relationship in a M1 pyramidal neuron, showing a decreased evoked firing rate during STN-DBS (2-way ANOVA followed by Bonferroni-corrected post-hoc tests, ***: p<0.001). STN-DBS increases the threshold (Wilcoxon signed-rank test, n=17, p=0.031), without affecting the gain of the *f-I* curve (paired t-test, p=0.17). (F) Single-shock STN stimulation induces a pause in M1 pyramidal evoked firing. Top: representative trial. Middle: raster plot of current-evoked activity for 10 trials in the same neuron. Bottom: the interval between the STN stimulation and the next evoked spike is longer than between the STN stimulation and the previous evoked spike (paired t-test, n=6 neurons, p=0.0057). (G) Top: a single-shock STN stimulation applied during hyperpolarizing or depolarizing current steps (-1.8 to +1nA in 0.2nA increments, mean of 10 trials) elicits a depolarizing or hyperpolarizing evoked potential. Bottom: voltage-dependency, displaying a reversion of the evoked potential around -71mV (inset) (n=10). DBS artifacts were removed in (D) and (E) for clarity.

In M1, GABAergic inhibition is provided by local neuronal populations mainly composed of parvalbumin(PV)-and somatostatin(SST)-expressing interneurons *(19).* To identify the cell-type specific populations recruited by DBS, we used genetically modified mice expressing channelrhodopsin (ChR2) in either PV *(Pv::ChR2* mice) or SST *(Sst::ChR2* mice) interneurons (Fig. 2 and Fig. S2). We verified that ChR2 was effectively expressed in targeted interneuronal populations and noted minimal ectopic expression (Fig. S2). In mice, stereotaxic targeting of the STN is challenging because of its small volume and inter-individual variability. To ensure the proper placement of the stimulation electrode, we first electrophysiologically localized the STN based on its typical firing activity, and in a second step implanted the DBS electrode at the same coordinates (Fig. S3). We then performed juxtacellular recordings of M1 neurons in *Pv::ChR2*and *Sst::ChR2* mice, allowing us to monitor DBS-evoked responses in opto-identified neuronal subpopulations (Fig. 2A, B and Fig. S3). Pyramidal, PV and SST cells were identified by their responses to light (or lack thereof) and their waveforms, as well as by juxtacellular labeling with neurobiotin for immunohistochemical and morphological identification (Fig. 2B and Fig. S3). Most (75%) of the recorded neurons were located in M1 layer V (Fig. S3). Consistent with previous *in vivo* recordings in anaesthetized mice *(20),* SST interneurons exhibited a lower spontaneous activity compared to PV cells. In line with the aforementioned results obtained in rats (Fig. 1), the firing activity of pyramidal cells was also decreased by STN-DBS in *Pv::ChR2*and *Sst::ChR2* mice (p=0.046, Fig. 2C, D and Fig. S3). We first investigated STN-DBS effect on PV interneurons. Indeed, they are the most likely candidates for recruitment by antidromic axonal reflex due to their inputs from cortico-STN pyramidal cell collaterals (21). Surprisingly, STN-DBS decreased the firing activity of PV interneurons (p=0.007). We next examined SST interneurons and observed that STN-DBS caused the opposite effect, increasing SST cell activity *(p=*0.0046) (Fig. 2C, D). These effects were robust across cell populations as 64% of the SST cells were excited whereas 70% of the PV cells were inhibited (Fig. 2E). Importantly, the kinetics of STN-DBS effect on pyramidal cells mirrored those of SST interneurons: we observed the same delay between the onset of STN-DBS and its effects on pyramidal and SST cells (Fig. 2D). Furthermore, pyramidal cell activity was strongly correlated with SST activity during and at the offset of STN-DBS (r=-0.89, p<0.001, compared with PV neurons: r=0.84, p<0.001), while their baseline activities were not correlated (r=;0.28, p=0.38). Therefore, it appears that STN-DBS efficiently drives activation of M1 SST interneurons, which could be responsible for the decreased activity of M1 pyramidal cells.

**Figure.2.**
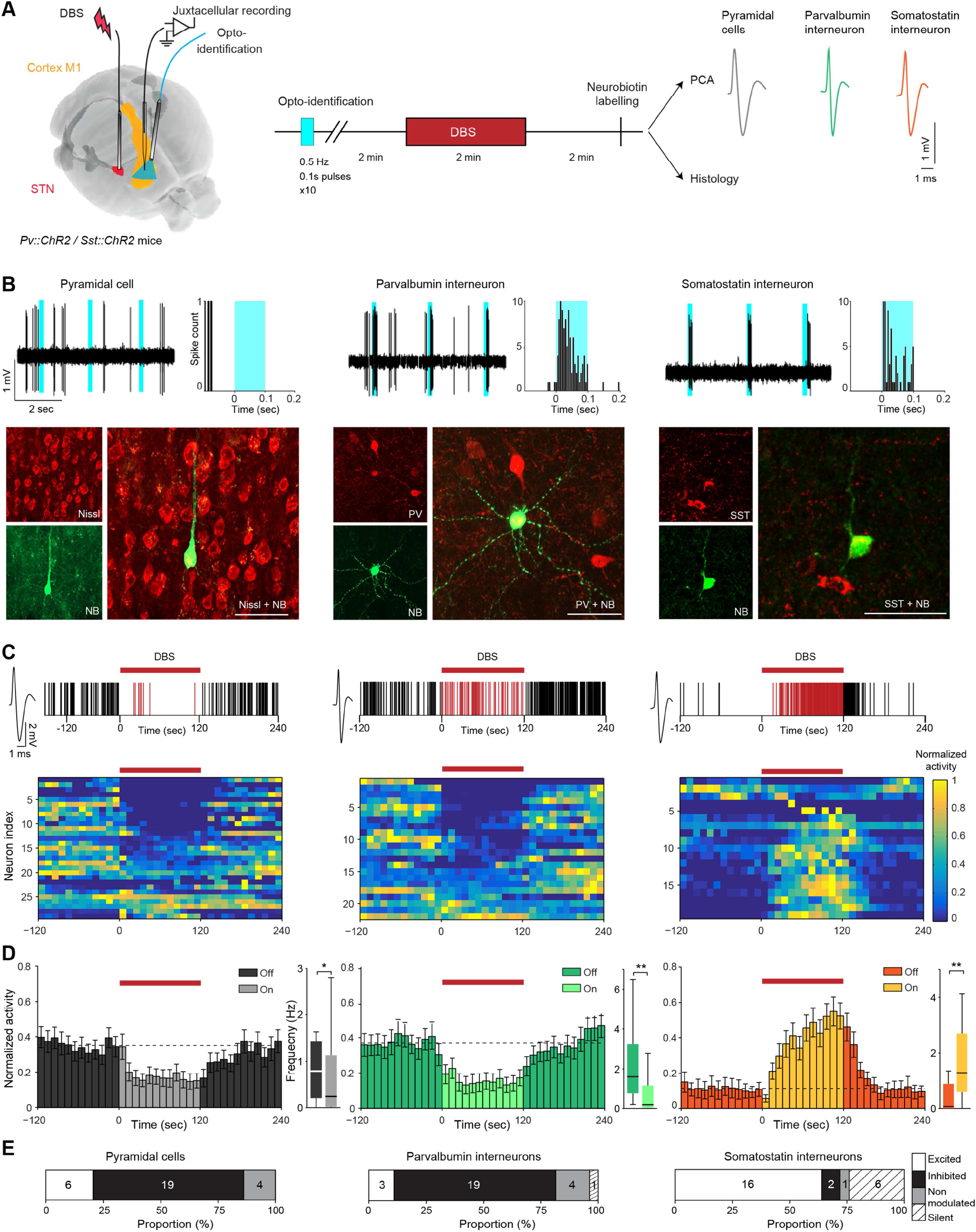
Somatostatin, but not parvalbumin, interneurons are recruited by STN-DBS. (A) *In vivo* experimental setup. A bipolar electrode is lowered into the STN and an optical fiber is placed over the M1, while recording from neurons in the M1. (B) Top: Electrophysiological traces of representative and identified pyramidal, SST and PV neurons recorded in the M1. SST and PV neurons are photo-activated by brief flashes of light (shown in light blue). Middle: Photomicrographs of juxtacellularly labeled and immuhistologically identified pyramidal, SST and PV neurons (bar scale=30pm). Bottom: Raster plots representing the activity of the neurons described above and their responses to a 2 minutes STN-DBS of 120pA intensity. Spikes occurring during the STN-DBS are represented in red. (C) Heatmaps of individual pyramidal, PV and SST neurons activity (normalized by the maximal firing rate of each neuron), before, during and after STN-DBS (10s bins). (D) Averaged time course of DBS-induced modulation of pyramidal, PV and SST neuron activity (mean±SEM). Boxplots indicate the firing rates before and during STN-DBS of pyramidal (Wilcoxon signed-rank test, p=0.046), PV (p=0.011) and SST (p=0.004) neurons. (E) Proportion of pyramidal, PV and SST neurons activated (white), inhibited (black), non-modulated (grey) during STN-DBS or silent (hatched) throughout the recording. The number of neurons is indicated for each category.

We thus hypothesized that cortical SST interneurons could constitute a target to efficiently mimic the effects of STN-DBS. Therefore, we tested whether the direct activation of SST interneurons would improve motor symptoms in freely moving parkinsonian mice. Unilaterally 6-OHDA-lesioned mice *(Sst::ChR2, Pv::ChR2* and wild-type mice) were implanted with an optical fiber in the ipsilateral M1 (Fig. 3A). We monitored the impact of opto-stimulation on the asymmetrical locomotor behavior induced by unilateral 6-OHDA-lesioning in two different tasks: the open field (Fig. 3B) and the cross-maze (Fig. 3C). In the open field, opto-activation of SST or PV interneurons decreased spontaneous ipsilateral rotations (p=0.0039 and p=0.020), whereas no effect was observed in wild-type mice (p=0.81) (Fig. 3B). This decrease in asymmetrical behavior did not result from a decreased locomotor activity, which remained unaffected by opto-activation of SST or PV cells (p=0.39 and p=0.11) (Fig. 3B and Fig. S4). In the cross-maze task, 6-OHDA-lesioned *Sst::ChR2, Pv::ChR2* and wild-type mice exploring the maze without opto-activation exhibited a strong bias towards ipsilateral turns (Fig. 3C) as expected for hemi-parkinsonian rodents, while sham-mice displayed random spontaneous alternance (Fig. S4B). In 6-OHDA-lesioned mice, only opto-activation of SST interneurons led to a decrease in the ipsilateral bias (p=0.0067) and an increase in the straight choice (p=0.0084) (Fig. 3C). Indeed, opto-stimulation in hemi-parkinsonian *Pv::ChR2* or wild-type mice failed to attenuate the ipsilateral bias (p=0.23 and p=0.93) (Fig. 3C). Interestingly, opto-activation of PV or SST interneurons in sham-mice did not induce an asymmetrical behavior contralateral to the opto-activation neither in the open field nor in the cross-maze task (Fig. S4B). This indicates that the reduced asymmetry observed upon SST opto-activation in 6-OHDA-lesioned mice is due to an improvement of pathological symptoms rather than a generic effect of unilateral opto-activation of cortical interneurons. Therefore, specific opto-activation of cortical SST interneurons alleviates the parkinsonian motor symptoms induced by a unilateral-lesion in mice.

**Figure.3.**
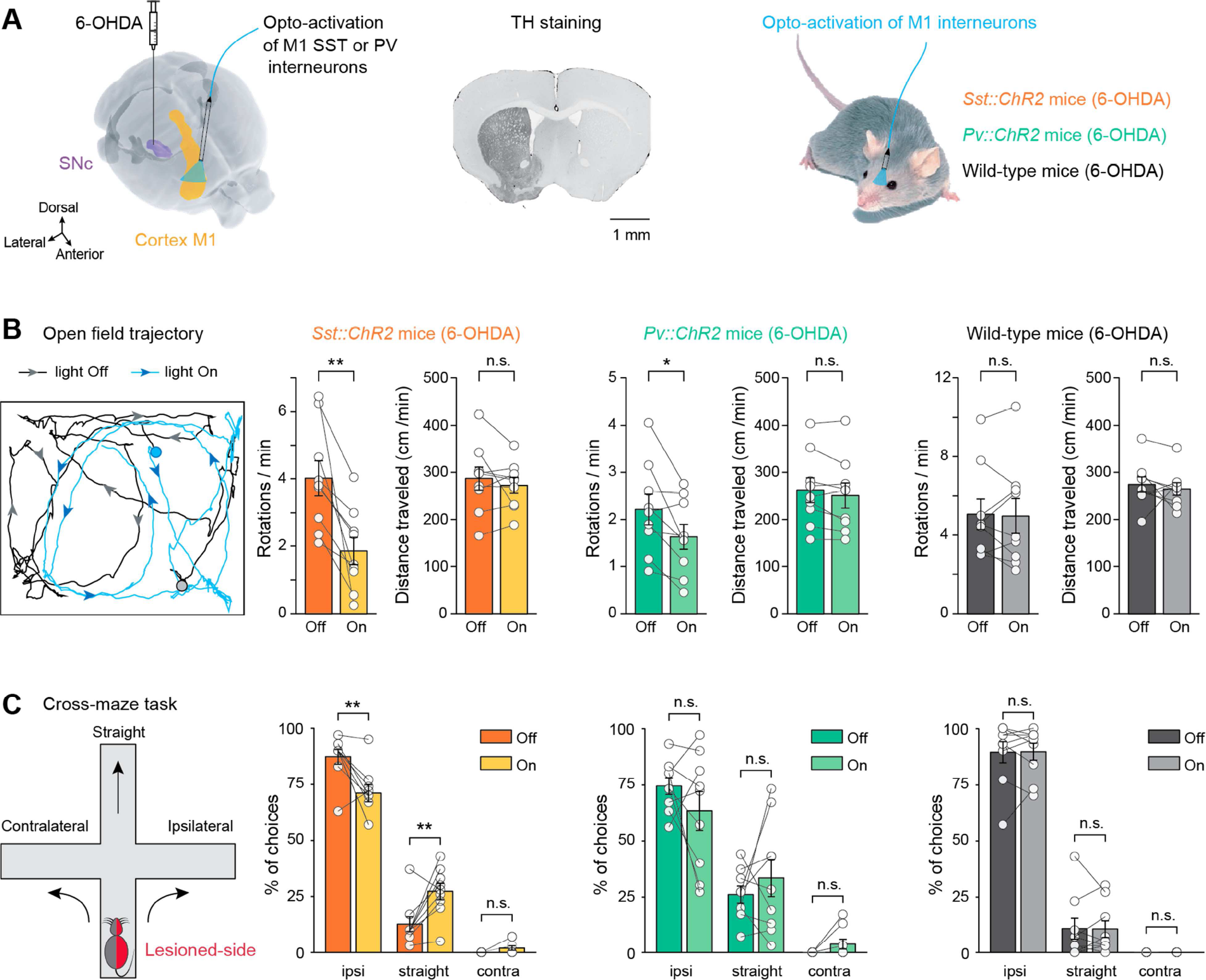
Opto-activation of M1 SST interneurons alleviates parkinsonian symptoms. (A) Experimental design: unilaterally 6-OHDA-lesioned (example TH immunostaining) *Sst::ChR2, Pv::ChR2* and wild-type mice *(n=9* in each group) were implanted with an optic fiber in M1 ipsilateral to the lesion. (B) Rotational behavior was quantified in an open field, in the presence or absence of light (left; 30s of trajectory shown in both conditions in a *Sst::ChR2*mouse. Circles and arrows indicate the starting point and direction of the animal). The number of rotations ipsilateral to the 6-OHDA lesion was significantly decreased during light-stimulation in both *Sst::ChR2* (p=0.0039, t-test) and *Pv::ChR2* (p=0.020, t-test) mice, but not in non-opsin expressing wild-type mice (p=0.81, t-test). Light stimulation did not affect the distance traveled (paired t-test, *Sst::ChR2* p=0.39; *Pv::ChR2* p=0.11; wild-type : p=0.56). (C) In the cross-maze test, ipsilateral turn preference was significantly decreased in favor of straight turns during light stimulations in *Sst::ChR2* mice (paired t-test, ipsi turns p=0.0067 and straight turns p=0.0084), but not in *Pv::ChR2* (p=0.84 andp=0.56) and wild-type (p=0.93 andp=0.93) mice.

To better understand the beneficial effects of STN-DBS and SST opto-activation, we theoretically tested the information processing capabilities of cortical networks in control and parkinsonian conditions under STN-DBS, PV or SST opto-activation. We built a simplified spiking neural network model of pyramidal cells, PV and SST interneurons (Fig. 4A; Table S1). In layer V motor cortex, both PV and SST interneurons inhibit pyramidal cells, but only PV cells receive excitatory feedback *(21)* (Fig. 4A). Hyperexcitability of pyramidal cells in the parkinsonian condition was taken into account by decreasing their firing threshold *(10).* STN-DBS was modeled as an excitatory input to all three populations. Consistent with our experimental findings, we observed a decrease in the firing rate of both pyramidal and PV cells while SST interneurons were activated at all tested frequencies and the impact was stronger with increasing stimulation frequency (p<0.001; Fig. 4B). More precisely, we observed that the decrease in pyramidal cell activity linearly scaled with the overall amount of current injected in the network. This phenomenon remained robust to changes in the parameters of 130 Hz DBS stimulation (pulse duration and amplitude) (Fig. S5). We next evaluated the relative contributions of PV and SST interneurons for driving changes in pyramidal cell activity under STN-DBS. We found a larger decrease of pyramidal cell activity in response to stimulations of SST compared to PV interneurons (Fig. 4C and Fig. S5), in agreement with experiments (Fig. 2). In particular, no decrease in pyramidal cell activity was observed in the absence of DBS-induced current on SST interneurons (Fig. 4C). We further simulated the impact of PV or SST opto-activation. As experimentally observed (Fig. S6), our model shows a remarkable inhibition of pyramidal cell activity for stimulations of either population at 67 and 130 Hz (Fig. 4D and E), yielding a sparse residual activity of pyramidal cells.

**Figure.4.**
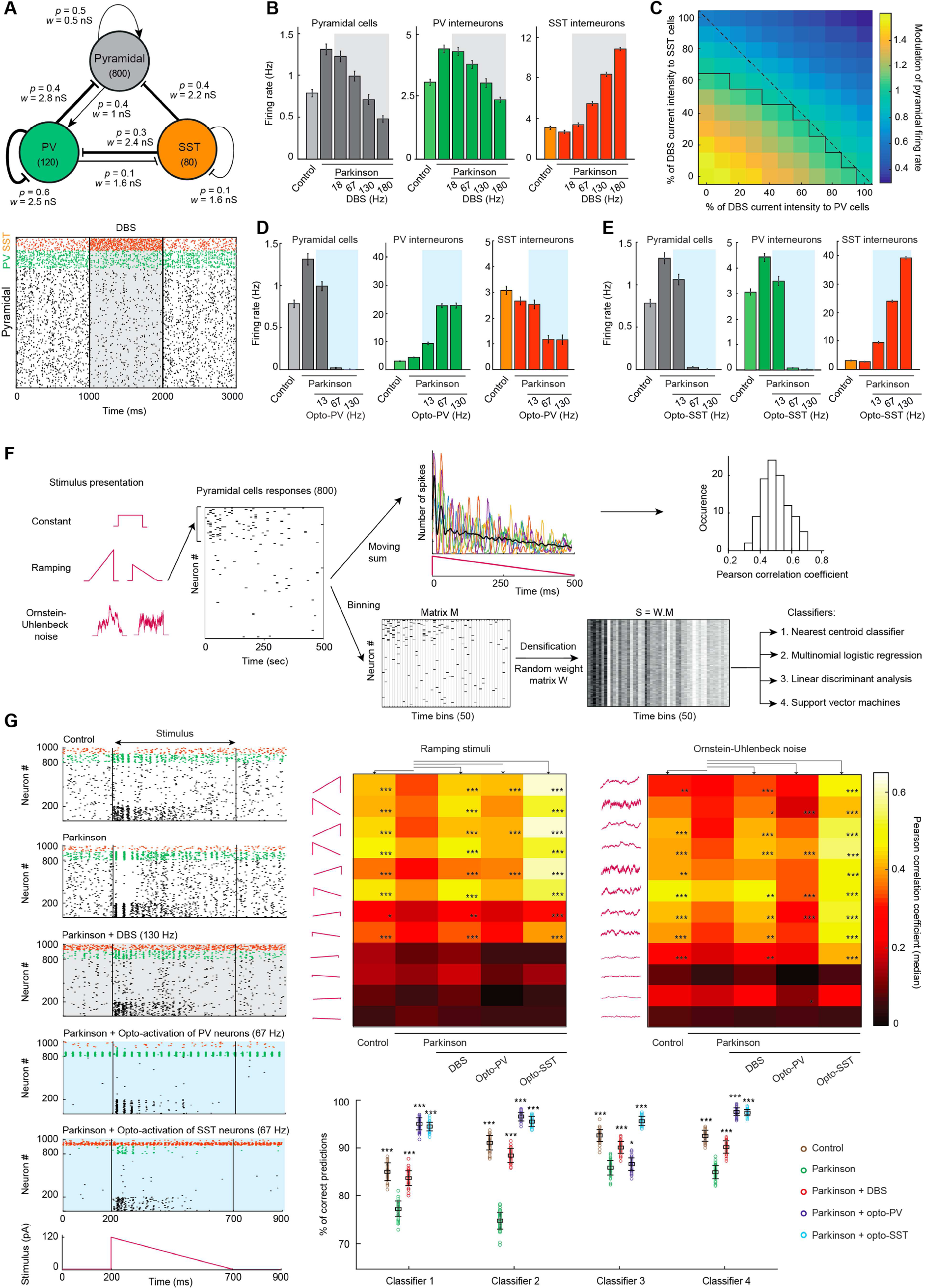
Theoretical impact of STN-DBS and opto-activation of PV and SST interneurons on cortical information processing. (A) Model of layer V motor cortex and network responses. *Top.* Network architecture: 800 pyramidal cells, 120 PV and 80 SST interneurons, with population-specific connection probability *p* and synaptic weight *w* (arrow: excitatory, bar: inhibitory). Raster plot before, during and after STN-DBS. (B) Average firing rate (±SD) for each population across different conditions and for multiple DBS frequencies (current amplitude: 120pA). All control and DBS conditions were significantly different from the parkinsonian condition (One-way ANOVA, p<0.001). (C) Average firing rate of pyramidal cells relative to the parkinsonian condition as a function of the percentage of STN-DBS current intensity received by SST and PV interneurons (maximal intensity: 120pA; frequency: 130Hz). 50% of pyramidal cells receive STN-DBS current. Black line: isocline corresponding to a ratio of 1. (D) and (E) Average firing rates (±SD) for all three populations under opto-activation of PV and SST interneurons, respectively. All firing rates during opto-activation were significantly different from those observed in parkinsonian conditions (One-way ANOVA and Tukey-Kramer *post-hoc* tests, p<0.001). (F) Procedure for calculating Pearson correlations between network responses and time-varying stimuli and training the classifiers. (G) *Left.* Raster plots of network responses to a ramp. *Center,*Heatmaps of median Pearson correlation coefficients for ramping stimuli and stochastic inputs. Significant differences with the parkinsonian condition are indicated (Kruskal-Wallis; *: p<0.05; **: p<0.01; ***: p<0.001) *Bottom.* Accuracy (±SD) of four different classifiers (One-way ANOVA and Tukey-Kramer *post-hoc* tests, *: p<0.05; ***: p<0.001).

We next asked how the STN-DBS-triggered pyramidal cell activity reduction could account for the improved motor function. To address this question, we studied the capacity of the network to transmit information, in both control and parkinsonian conditions, and during the application of STN-DBS, PV or SST opto-activation. The impact of the modifications of the network activity on information transmission in these conditions is not heuristically obvious: in Parkinson’s disease, the increased excitability of pyramidal cells would make the system more responsive to stimuli, but the higher spontaneous activity may interfere with stimulus-evoked activity. In contrast, the lower spontaneous activity induced by DBS and DBS-guided opto-activation may avoid signal degradation, but could bring the system to a less responsive state. We first tested whether pyramidal cells conserve the capacity to generate specific patterns of activity in response to stimuli despite sparse activity under DBS or opto-activation. We applied 28 inputs with different motifs (deterministic constant and ramping stimuli, and stochastic Ornstein-Uhlenbeck processes; Table S2) and intensities to subsets of pyramidal cells (Fig. 4F and G), and systematically evaluated the information conveyed by the network. We found that for medium to high amplitude stimuli, pyramidal cell responses show lower levels of correlation with the stimulus in parkinsonian condition compared to control. Both STN-DBS and SST opto-activation, but rarely PV opto-activation, increased the correlation coefficients when compared to parkinsonian condition (Fig. 4G and Fig. S7). Low amplitude stimuli yielded weak correlations that were not significantly different across conditions and thus did not confer any advantage to the parkinsonian condition. Furthermore, patterns of activity in the motor cortex could determine how downstream areas, such as the striatum, can decode the transmitted signal. We used four machine-learning decoding algorithms (nearest centroid classifier, multinomial logistic regression, linear discriminant analysis and support vector machines; Table S3) to discriminate network responses to 20 different stimuli (Fig. 4E). Network responses were densified to mimic highly convergent cortico-striatal information transfer. Classifier accuracy was higher in control and DBS relative to parkinsonian condition (for all classifiers: p<0.001), and opto-activation similarly improved the classification accuracy compared to parkinsonian condition (for linear discriminant analysis: p<0.05 for PV opto-activation and p<0.001 for SST opto-activation; for the three others classifiers: p<0.001 for PV and SST opto-activation; Fig. 4G). Thus, our modeling results indicate that reducing pyramidal cell hyperexcitability enables the cortical network to efficiently encode stimulus content and may facilitate the extraction of this information by downstream areas.

STN-DBS has been widely described as acting on distinct basal ganglia nuclei *(4-6)* and recent studies also point towards a significant impact of STN-DBS at the cortical level *(7, 8, 11-17).* A better understanding of the cortical effects of STN-DBS might allow transposing its remarkable efficacy to less invasive therapeutic strategies, thus making it available for a larger number of parkinsonian patients. Here, we show that DBS counteracts the hyperactivity of M1 pyramidal cells in parkinsonian rodents, in agreement with data collected from patients *(12-17).* Our *in vivo*intracellular recordings indicate that DBS activates cortical GABAergic networks. Using opto-identification of cortical interneurons, we report that STN-DBS leads to a dualistic action on cortical GABA interneurons, notably activation of SST and inhibition of PV interneurons. Furthermore, opto-activation of cortical interneurons alleviates motor symptoms in parkinsonian rodents. Interestingly, our theoretical modeling reveals that STN-DBS and interneuron opto-activation counteract the cortical hyperexcitability, thus restoring information processing capacities of the cortical network. The asymmetrical connectivity of SST interneurons with layer V pyramidal cells (21) implemented in our model accounts for the higher efficiency of SST compared to PV opto-activation (Fig. 3 and 4). Antidromic activation of the cortico-subthalamic axons, orthodromic activation of subthalamo-cortical fibers and/or basal ganglia-thalamo-cortical loops could mediate SST recruitment by STN-DBS *(4, 5,* 11). The direct activation of SST interneurons by antidromic axonal reflex is unlikely to occur since cortico-STN pyramidal cells lack collaterals to SST interneurons (21); the delayed activation of SST interneurons by STN-DBS further argues against this hypothesis. This implies that the evoked short latency inhibitory potential (Fig. 1G and Fig. S1B) is unlikely due to SST recruitment. The orthodromic STN-cortex pathway could account for the SST recruitment, since it mainly targets superficial cortical layers (*22*), where pyramidal cells directly connect layer V SST interneurons (*21*). The involvement of the basal ganglia-thalamo-cortical pathways might explain the delay in SST interneuron recruitment, in line with the slow but facilitating kinetics of thalamic inputs to cortical SST neurons observed in the somatosensory cortex *(23, 24).* The recruitment of cortical SST interneurons most likely relies on the combination of direct and indirect effects of steady-state DBS on cortico-basal-ganglia-thalamic networks.

Recent studies highlight the key roles operated by SST interneurons in cortical information processing *(25-27).* Their impact, which results from their specific input and output connections, differs depending on SST subpopulation across cortical layers (28). Additional work is needed to determine if DBS differentially recruits distinct cortical SST subpopulations, in order to better refine therapeutic targets. The beneficial effect of the activation of SST interneurons as observed in this study, and other pathological contexts *(29),* highlights the hyperactivity of pyramidal neurons as a neuropathological marker, and cortical inhibitory interneurons as a promising therapeutic target. Besides SST, other interneuronal populations might constitute alternative targets. Indeed, here we show that although PV interneurons are inhibited by STN-DBS, their specific activation partially alleviates motor symptoms in parkinsonian mice, and restores cortical processing, albeit less efficiently than SST opto-activation. Therefore, increasing the inhibitory drive into the motor cortex could represent a useful strategy to alleviate motor impairments. Targeting combinations of interneurons might increase the therapeutic benefits of our strategy. While there is an emerging therapeutic potential of optogenetics *(30-35),* its medical application is still in its infancy, with the first human clinical trial underway to treat retinitis pigmentosa *(clinicaltrials.gov,* NCT03326336 and NCT02556736). More work is necessary to fully explore and optimize safe CNS transfection and light delivery in humans *(34-36).* Nevertheless, targeting cortical GABAergic networks with pharmacology or non-invasive brain stimulation (36) such as trans-cranial stimulation (37) could provide less invasive strategies than STN-DBS, thus benefiting to a larger population of parkinsonian patients.

## Acknowledgments

We thank the L.V. lab members, Christian Giaume, Nicolas Gervasi, Pierre Magistretti and Yulia Worbe for helpful suggestions and critical comments. We thank Sylvie Perez, Alessandra Romei, Farah Hadj-Idris, Angèle Roudeau and Charlotte Branco for technical assistance for 6-OHDA lesioned-rodents and immunohistochemistry;

## Authors contributions

Conceptualization: L.V., B.D. and J.T.; B.D. performed *in vivo* intracellular electrophysiological experiments; S.V., B.D., AAA and WD performed *in vivo* extracellular electrophysiological experiments; S.V. performed *in vivo* juxtacellular electrophysiological, opto-identification and immunohistochemistry experiments; MV and GG performed *in vivo* optogenetic, behavioral and immunohistochemistry experiments; M.V., B.D., S.V., C.P., G.G. and L.V. performed analysis; B.D., G.G. and M.V. performed 6-OHDA lesions; J.T. and C.P. carried out the conception and the design of the mathematical model; C.P. performed the acquisition and analysis of data from the mathematical model; L.V., B.D., J.T., M.V., S.V., C.P. wrote the manuscript and all authors have edited and corrected the manuscript; Funding acquisition: B.D. and L.V.; L.V. supervised the whole study.

## Data and materials availability

All data is available in the main text or the supplementary materials. Moreover, all data, code, and materials used in the analysis is available in some form to any researcher for purposes of reproducing or extending the analysis.

## List of Supplementary Materials

Materials and Methods

Supplementary Figures S1 to S7

Supplementary Tables S1 to S3

References only cited in the SM: *38-57.*

## Materials and Methods

### Animals

All experiments were performed in accordance with the guidelines of the local animal welfare committee (Center for Interdisciplinary Research in Biology Ethics Committee) and the EU (directive 2010/63/EU). Every precaution was taken to minimize stress and the number of animals used in each series of experiments. Animals were housed in standard 12-hour light/dark cycles and food and water were available *ad libitum.*

Adult male OFA rats (n=31, 175-200 gr) (Charles River, L’Arbresle, France) were used for anesthetized intracellular and unitary extracellular recordings. Adult mice of both sexes (n=101, 2-12 months) from 3 mouse strains were used for anesthetized extracellular unitary recordings combined with opto-tagging, chronic freely moving behavior, and immunostaining: wild type C57Bl6 (Charles River), and transgenic hybrid *Pv::ChR2* and *Sst::ChR2.* The hybrid transgenic mice were heterozygous for both genes, obtained by mating a cre-driver transgenic line ensuring specific expression of the cre recombinase in PV or SST neurons (PVcre: Jackson Laboratory stock # 008069 or SSTcre: Jackson Laboratory stock #013044, Charles River) with a transgenic reporter line containing the light-activated channelrhodopsin-2(H134R) in a cre-dependent construct (ChR2: Jackson Laboratory stock #012569, Charles River).

### 6-OHDA lesions

Rats were anesthetized with an initial dose of sodium pentobarbital (30 mg/kg, i.p.), and adjusted to a surgical plane with injections of ketamine (27.5 mg/kg, i.m.; Imalgene, Merial, Lyon, France) repeated as needed. Mice were anesthetized with an i.p. injection of a mix of ketamine (67 mg/kg) and xylazine (13 mg/kg, Rompun, Bayer, Puteaux, France). Lidocaine (Xylovet, CEVA, Paris, France) was applied cutaneously. All animals received a bolus of desipramine (Tocris, Bio-Techne, Lille, France) dissolved in saline (rats: 25 mg/kg, i.p.; mice : 20 mg/kg, i.p.) 30 minutes before the injection of 6-hydroxydopamine (6-OHDA; Sigma, Steinheim, Germany) to prevent neurotoxin-induced damage of noradrenergic neurons. A single stereotaxic injection of 6-OHDA was performed. For rats, 7 μL of 6-OHDA (2.5 mg/ml in vehicle), or 7 μL of vehicle (saline containing 0.01% w/v ascorbic acid, Sigma) was injected at 16 μL/h through a steel cannula (0.25 mm outer diameter) attached to a 10 μL Hamilton microsyringe (Cole-Parmer, London, UK) controlled by an electric pump (KDS100; KD Scientific, Holliston, MA, USA) in the substantia nigra *pars compacta* (coordinates: 3.7 mm anterior to the interaural line, 2.1 mm laterally, 7.55 mm depth from the cortical surface). For mice, 0.5 μL of 6-OHDA (6 mg/ml in saline containing 0.02% w/v ascorbic acid) or 0.5 μL of vehicle (saline containing 0.02% w/v ascorbic acid) was injected at 3 μL/hour through a 33ga needle (NRS adapter kit) in the medial forebrain bundle (coordinates from bregma: 1.2 mm posterior, 1.2 mm lateral, 4.9 mm depth). Mice were placed in a thermostatic chamber at 32°C, their weight was monitored daily and they received s.c. injections of 0.8 mL glucose (50 mg/mL, i.p.) and/or saline as needed until recovery. Mice who failed to display spontaneous ipsilateral rotations in the 2 weeks following surgery were excluded from the study.

### In vivo intracellular recordings

After three weeks of recovery, 6-OHDA-lesioned and sham-lesioned rats were anesthetized in an induction chamber containing isoflurane 4.5% (AErrane®, Baxter S.A., Lessines, Belgium). A cannula was inserted into the trachea and isoflurane was delivered at 3-3.5%. Small craniotomies were drilled above the motor cortex (12.5 mm anterior to interaural line; 3.8 mm lateral) and STN (5.2 mm anterior; 2.5 mm lateral), and the dura matter was removed under isoflurane anesthesia (Univentor 400, Univentor Limited, Zejtun, Malta). For electrophysiological recordings, rats were maintained in a narcotized and sedated state by injections of fentanyl (4 μg/kg, i.p.; Janssen-Cilag, Issy-Les-Moulineaux, France) repeated every 20-30 min. To obtain long-lasting stable intracellular recordings, rats were immobilized with gallamine triethiodide (40 mg, i.m., every 2 h; Specia, Paris, France) and artificially ventilated (UMV-03, UNO, Zevenaar, Netherlands). Body temperature was maintained at 36.5°C with a homeothermic blanket (Harvard Apparatus, Kent, UK). At the end of the experiments, animals received a lethal dose of sodium pentobarbital (150 mg/kg, i.p.) and the brain was removed for histological processing.

Intracellular recordings were performed using glass micropipettes filled with 2 M potassium acetate (40-70 MQ). Intracellular recordings were obtained using the active bridge mode of an Axoclamp 2B amplifier (Molecular Devices, Union City, CA). Data were sampled on-line on a computer connected to a CED 1401 interface using the Spike2 data acquisition program (Cambridge Electronic Design, Cambridge, UK) with a sampling rate of 25 kHz for off-line analysis using Spike 2.

The STN ipsilateral to the recorded pyramidal cells was stimulated with a bipolar coaxial stainless steel electrode (SNE-100; Rhodes Medical Instruments, Woodlands Hill, CA) at 8.1 mm depth (38). Electrical stimulation consisted in either single-shock STN stimulations (60 μs, 2-4 V) or DBS (continuous stimulation at 130 Hz with the same parameters), using a stimulus isolator (DS2A, Digitimer, WelWyn Garden City, UK) driven by a pulse stimulator (Pulsemaster A300, WPI, Hitchin, UK).

Spontaneous activities of cortical neurons were recorded for at least 150 seconds before, during and after STN DBS. The pyramidal cell transfer function was quantified in the presence and absence of DBS as the relationship between the intensity of intracellularly injected currents and firing responses *(f—I* relationships). The firing rate was measured in response to depolarizing current pulses of increasing intensity (200 ms, 0-1.2 nA in 0.2 nA steps, with an inter-stimulus interval of 1.25-2.25 s). Each current intensity was applied at least 10 times in each condition (with or without DBS), and we averaged the firing rates obtained for each trial. Linear regressions were applied to *f—I* curves to determine the threshold current for AP generation *(I****threshoid***), extrapolated as the x-intercept of the linear fit, and the neuronal gain, defined as the slope of the *f-I* curve. The voltage-dependency was measured in response to a single-shock STN stimulation triggered during hyperpolarizing and depolarizing current pulses (0.3-1 s, -1.8 to +1 nA in 0.2 nA steps; STN stimulation 50-200 ms after current onset). For each current intensity, the membrane potential was averaged over 10 trials, and the latency and amplitude of the evoked response were measured. For each neuron, the voltage-dependency curve was first smoothed (lowess method), and the reversal potential was linearly interpolated as the y-axis intercept, using built-in Matlab functions (The Mathworks, Natick, MA).

### *In vivo* **juxtacellular recordings**

*Rats.* Unilaterally-lesioned or sham-lesioned rats were anesthetized with chloral hydrate (400 mg/kg, i.p., supplemented by continuous intraperitoneal injection delivered at a rate of 60 mg.kg^-1^.h^-1^ using a peristaltic pump). Small craniotomies were drilled above the motor cortex (12.5 mm anterior to the interaural line; 3.8 mm lateral) and STN (5.2 mm anterior; 2.5 mm lateral), and the dura was carefully removed. Body temperature was maintained at 36.5°C with a homeothermic blanket (Harvard Apparatus). At the end of the experiments, animals received a lethal dose of sodium pentobarbital (150 mg/kg, i.p.) and the brain was removed for histological processing. Single-unit activity of motor cortex cells was recorded extracellularly using glass micropipettes (10-15MQ) filled with a 1.0 M sodium chloride solution. Single neuron action potentials were recorded using the active bridge mode of a Axoclamp-2A amplifier (Molecular Devices), amplified, and filtered with an AC/DC amplifier (DAM 50; WPI) and displayed on a digital oscilloscope (TD 3014 B; Tektronix, Courtaboeuf, France). Spikes were discriminated from noise and stimulation artefacts on the basis of their amplitude, using the gate function (double threshold) of the discriminator (121 window discriminator; WPI) and sampled online on a computer connected to a CED 1401 interface using the Spike2 data acquisition program (Cambridge Electronic Design) at 10 kHz. The STN ipsilateral to the recorded pyramidal cells was stimulated with a bipolar coaxial stainless steel electrode (SNE-100; Rhodes Medical Instruments) at 8.1 mm depth, using the same DBS stimulation parameters as used in intracellular recordings. Spontaneous activities of cortical neurons were recorded for at least 120 seconds before, during and after DBS. Spikes were manually checked offline to ensure the correct discrimination from DBS artefacts.

*Mice.* Mice were anesthetized using urethane (1.5 g/kg i.p.) with supplementary doses as necessary until absence of reflexes following a hind paw pinch. Body temperature was maintained at 37 °C. A small craniotomy was made over the STN (1.8mm posterior to bregma and 1.5mm lateral to the midline) and the dura mater was removed. Recording electrodes were made from borosilicate glass capillaries (Harvard Apparatus), pulled with a vertical electrode puller (Narishige, London, UK) and tips were broken under microscope control (tip diameter ~1.5μm). Electrodes contained saline solution (0.5M NaCl) and Neurobiotin (1.5%) yielding impedances of 15-25 MQ. A reference electrode was placed subcutaneously and extracellular recordings of STN action potentials were performed by lowering the electrode with a micro drive to a depth of ~4.3mm. Electrode signals were amplified using an Axoclamp-2B amplifier (Axon Instruments), audio monitored (A.M. Systems, Phymep, Paris, France), digitized at 25 kHz and stored in Spike2 (Cambridge Electronic Design). STN position was identified with the known characteristic discharges of STN neurons in urethane anesthesia *(39).* Bursts in STN neurons were detected using the Poisson surprise method *(40).* To confirm STN position, some STN neurons were electroporated by application of positive current steps and expulsion of neurobiotin from the electrode, which filled the neurons, as previously described (47). Once the STN was located, electrodes were elevated out of the brain and a concentric bipolar electrode (CBDSF75, FHC, Bowdoin, ME) was targeted into the STN using the same coordinates. For optogenetic identification of neuronal populations, a craniotomy was made over the motor cortex ipsilateral to the STN (coordinates: 1.5mm anterior of bregma and 1.5mm lateral to the midline). An optical fiber mounted to a blue light source (Plexbright 465nm, Plexon, Dallas, TX) was placed on the cortical surface. Glass electrodes were lowered through the cortex while flashing brief pulses of light (100 ms, 0.2 Hz, 10 mW). After opto-identification, neurons were allowed to recover from light stimulation and spontaneous activity was defined on the basis of 2 min of stable recording before launching the DBS protocol. Electrical DBS consisted of 60μs pulses at 130 Hz during 2 min at 120 μA using a stimulus isolator (IsoFlex, A.M.P.I, Jerusalem, Israel) driven by a pulse stimulator (Master-8, A.M.P.I). Finally, neurons underwent a series of 1 min opto-activations at 13, 67 and 130 Hz (3 ms pulse), with at least 1 min of recovery between each series. At the end of the experiment, neurons were juxtacellularly labeled when possible, and electrolytic lesions of the STN were performed (30μA DC for 30 sec) for*posthoc* confirmation of appropriate targeting of the bipolar electrode to the STN. All data obtained from animals with lesions outside the STN were excluded. Spikes were manually checked offline to ensure the correct discrimination from DBS artefacts. Because *Pv::ChR2* and *Sst::ChR2* mouse strains can display a small proportion of off-target expression of ChR2 (fig S2), we used cluster analysis to combine opto-activation and waveform characteristics in order to better identify recorded neurons (fig S3). For each cortical neuron, the waveform was filtered, averaged, and normalized by their peak amplitude. Principal component analysis (Matlab) was then applied to waveform characteristic features (coordinates of the trough and of the second peak in the normalized waveforms) and opto-activation success rate. Neurons were then clustered using hierarchical and k-means clustering algorithms (Matlab), testing all combinations of the first 2, to all principal components, with 2 to 7 target clusters. The resulting cluster silhouettes were ranked and the best score was selected (mean silhouette =0.74, with 2 principal components, 2 clusters, k-means algorithm), resulting in two clearly separated clusters. Morphological and immunohistochemical identity of labeled neurons (n=18) allowed us to identify the two clusters as pyramidal cells and interneurons. Two cells where the morphological and clustered identity did not match were excluded from the analysis.

### Behavior

*Optic fiber implantation.* After recovery from the 6-OHDA injection, mice were anesthetized in an isoflurane chamber induction followed by ketamine:xylazine mix (67:13 mg/kg i.p). Additional isoflurane was adjusted (0.5-1%) throughout the procedure to maintain a deep anesthesia. Temperature was maintained constant using a heating pad (Gaymar T/Pump, WPI). Two screws were inserted symmetrically in parietal skull plates for securing the implant. An optic fiber cannula (200 μm core, 0.39 NA, CFML 12U-20, Thorlabs, Maisons-Laffitte, France) was implanted in the motor cortex (1.2 mm anterior, 1.5 mm lateral to bregma at 0.2-0.4 mm below the cortical surface). After closing the craniotomy with silicon (Kwik-Cast, WPI), the implant was cemented to the skull and protected by a coppermesh circlet embedded in the cement. Mice received 0.8 mL of saline (s.c.) after the procedure, and were monitored postoperatively until recovery.

*Light stimulation.* Blue light (475 nm) was delivered from a laser diode light source (FLS-475, DIPSI, Cancale, France) through a rotary joint (FRJ_1×2i_FC-2FC, Doric Lenses, Québec, Canada) connected to the mouse optic fiber cannula, controlled by a Master-9 (A.M.P.I.) delivering 3 ms light pulses at 67 Hz. The light intensity (10 mW max at the tip) was adjusted before each recording using an optometer (PM100D + S130C, Thorlabs). The light stimulation and the synchronized video, including animal tracking, were recorded using a KJE-1001 recording system (Amplipex, Szeged, Hungary).

*Behavioral tasks. Open field*: mice were placed in a rectangular arena (40 x 25 cm, with walls of 20 cm height). After a habituation period of 30 sec, the spontaneous locomotor activity and rotations were monitored, alternating 1 minute with the light off and 1 minute with the light on (3 ms pulses at 67 Hz), for a total duration of 10 min. The distance travelled was evaluated with custom Matlab scripts from the Amplipex tracking data; the rotations were counted manually on the recorded video. *Cross-maze:* mice were placed in one arm of a cross-maze (arms: 36.5 x 7 cm, with walls of 10.5 cm height) and left free to explore the maze until they performed at least 30 turns (left, right or straight) with a 30 minute cut-off. Right turns corresponded to turns towards the ipsilateral side of the dopaminergic lesion *(42).* Two sessions were performed: a control session where the mouse was connected to the optic fiber, but with the light off, and a session with cortex light stimulation (3 ms pulses at 67 Hz) applied continuously during the whole session. Videos were analyzed offline, and the percentage of turn in each direction was calculated on the first 30 turns.

### Histology

*Assessment of 6-OHDA lesion.* Tyrosine hydroxylase (TH) immunohistochemistry was performed to assess the extent of the lesion. After fixation of the brain (4% w/v paraformaldehyde in phosphate buffer), coronal slices (50-80 μm thickness) of the striatum and SNc were cut with a cryotome (HM400 and KS34, Microm Microtech, Brignais, France) and processed for TH-immunostaining, either (i) with immunofluorescence: using a mouse anti-TH primary antibody (1/300-1/500, 24h at 4°C, MAB318, Merck Millipore, Fontenay-sous-Bois, France) and a donkey anti-mouse coupled to Cy5 fluorophore (1/500, 2h at RT, 715-175-151, Jackson ImmunoResearch, Interchim, Montlu?on, France), counterstained with DAPI (1/5000, 10 min at RT, Molecular Probes, Thermofisher, Illkirch, France); or (ii) with immunohistochemistry, after endogenous peroxidases inactivation, using a rabbit anti-TH (1/1000, 36h at 4°C, AB152, Merck Millipore), and biotinylated goat antibody against rabbit IgG (1/200, BA 1000, Vector Laboratories, CliniSciences, Nanterre, France), followed by revelation using an ABC kit (PK-6100, Vector Laboratories). Images were captured using an epifluorescence microscope (DMRB, Leica, Nanterre, France) or a stereozoom fluorescence microscope (Axiozoom, Zeiss, Marly le Roi, France), and analyzed using ImageJ software. *Efficacy and specificity of transgenic mouse lines.* Two mice of each transgenic hybrid strain *(Pv::ChR2* and *Sst::ChR2)* were euthanized by injection of a lethal dose of anesthetic (sodium pentobarbital, >150 mg/kg, i.p.), then transcardially perfused with saline followed by a fixative solution (4% w/v paraformaldehyde in phosphate buffer), and was then post-fixed overnight at 4°C. Coronal slices (30-50 μm) of the motor cortex were cut in PBS using a vibratome (HM650V, Microm Microtech) in PBS, and processed for immunostaining of either PV or SST, and YFP (ChR2 expression reporter). For SST / YFP immunostaining, an extra step of antigen retrieval was performed in citrate buffer for 15 min at 75°C. The following primary antibodies were used: rabbit anti-PV (1/2500, PV25, Swant, Marly, Switzerland), rat anti-SST (1/100, MAB354, Merck Millipore), chicken anti-GFP (1/1000, AB13970, Abcam, Paris, France), incubated 24h (PV/YFP) or 72h (SST/YFP) at 4°C; and secondary antibodies : donkey antirabbit coupled to Alexa647 (1/500, 711-605-152, Jackson ImmunoResearch), donkey anti-rat coupled to Alexa647 (1/500, 712-605-153, Jackson Immuno Research), and goat anti-chicken coupled to Alexa488 (1/500, 103-545-155, Jackson Immuno Research) incubated 1-2h at RT. Slices were observed and images were captured using a confocal microscope (SP5, Leica), and analyzed using ImageJ software.

*Revelation of juxtacellular labelling.* Following perfusion and brain extraction, 60 μm thick coronal sections were serially cut with a vibratome (HM650V, Microm Microtech). Floating sections were incubated (2 h) at room temperature in Alexa 488-conjugated streptavidin (1/250, S11223, Invitrogen, Thermofisher) in PBS containing 0.2% Triton X-100. Sections were mounted on glass slides for examination under an epifluorescence microscope (DMRB, Leica), and sections with the labeled neurons were dismounted from the slides for further revelation. Then, sections were processed for immunostaining of SST and PV as described above, and images were taken with a stereozoom fluorescence microscope (Axiozoom, Zeiss), and analyzed using ImageJ software.

### Statistics

Unless otherwise stated, normal data are displayed as mean±SEM, and differences between groups were assessed using unpaired or paired t-test. Non-normal data are presented as boxplots where the white line is the median, the box represents the 25% and 75 % quartiles (Q1 and Q3), and the whiskers represents 1.5* the interquartile range outside of the box range (i.e., Q1-1.5*(Q3-Q1) for the bottom whisker, and Q3+1.5*(Q3-Q1) for the top whisker), extended to the adjacent value (data points outside the whisker range are not shown for clarity of display, but are included in the statistical analysis), and differences between groups were assessed using Mann-Whitney U test (unpaired data) and Wilcoxon’s signed rank test (paired data). Normality of each dataset was tested using D’Agostino and Pearson’s test.

### Model

*Spiking network model.* We built a simplified spiking model of the layer 5 of motor cortex: the network consisted of 800 pyramidal cells, 120 PV and 80 SST interneurons, consistent with the 1:5 ratio between cortical excitatory and inhibitory cells and the larger proportion of PV ***vs.*** SST neurons in L5 (***43***). Neuron of index ***i*** was modeled as an adaptive exponential integrate-and-fire neuron (44): its activity depends upon the dynamics of a fast voltage variable v^i^ and a slow adaptation variable ***w***^i^, according to the following stochastic equations (1) and (2):

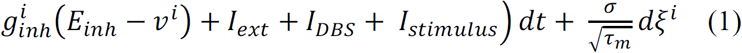

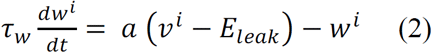

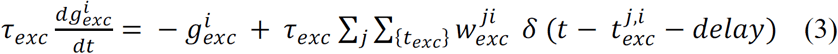

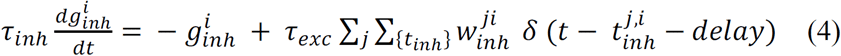

The value of the voltage ***v***^i^ is thus determined by intrinsic dynamics and input integration mechanisms. The intrinsic dynamics consists of a linear relaxation towards the resting state (with a time constant τ_m_ = C/g_leak_) and an exponential spiking current active once the spike threshold *Vthres* is reached, modeling the sharp onset of spike generation. The voltage also integrates excitatory/inhibitory synaptic inputs from the network as well as external stimulations: external constant currents I_ext_, specific stimuli *I_stimulus_*, DBS-induced currents *I_DBS_,* as well as noisy currents modeled as independent Gaussian white noise (5^i^). These stochastic inputs encompass the variety of sources of fluctuations of the voltage *(45),* whose amplitude is determined by the parameter σ. The voltage also experiences a negative feedback modeling adaptation; this phenomenon is taken into account through the slower variable *w*^i^, whose dynamics is linear in *v*^i^ and controlled by the parameter *a* determining the strength of the adaptation-voltage coupling.

A spike is triggered when the voltage approaches the threshold V_thres_, specific to each population. Upon firing, the variable *v*^i^ is reset to a fixed value, here equal to the leak reversal potential, *V_reset_* = *E*_leak_ = -60 mV (44), whereas *w*^i^ is increased by a fixed amount b, corresponding to the spike-triggered adaptation. The values of all intrinsic parameters are summarized in Table S1. We underline the following choices motivated by experimental data. First, pyramidal cells had a larger leak conductance *g*_leak_ and capacitance *C* compared to interneurons. The spike threshold of pyramidal cells was adjusted in parallel with the amount of constant external current *I*_ext_ received such that, in absence of stimulus, their firing rate remained near 1 Hz, consistent with experimental observations of sparse activity and low-firing rates in awake cortical areas *(46).*Secondly, PV neurons were modeled as fast-spiking interneurons, with a sharp spiking onset and no adaptation *(47)* whereas the profile of SST interneurons was characterized by a lower threshold for spike generation and the existence of spike-frequency adaptation (25). Adaptation parameters for pyramidal and SST neurons were in the range of those proposed by Brette and Gerstner (44) and Naud et al. *(48).* Finally, the parkinsonian model incorporated the hyperexcitability of pyramidal neurons through a decreased spiking threshold compared to the control condition *(10),* chosen to match the experimentally observed increase in firing rate.

Synaptic connections between neurons were randomly distributed, with a given connection probability *p* ranging from 0.1 to 0.6 and a fixed synaptic weight *w* (Fig. 4A). We opted for a simple conductance-based description of synaptic currents: the excitatory and inhibitory conductances *g*_exc_ and *g*_inh_ display a discrete jump following action potentials (arriving at times *t*_exc_ or *t*_inh_), after a delay of 1 ms, and decay exponentially with a time constant *τ*_exc_ = 3 ms and *τ*_inh_ = 5 ms (equations (3) and equations (4) respectively). The excitatory *E_exc_* and inhibitory *E_inh_* reversal potentials were set to 0 mV and -80 mV respectively *(49).*

The network connectivity was determined based on published experimental results *(21, 50-52)*and previous models of the motor cortex (53). In particular, the absence of connection from pyramidal cells to SST interneurons has been previously reported (21). When choosing the synaptic weights, we set lower conductances for excitatory synapses compared to inhibitory ones *(49, 54)* and respected the observation that PV interneurons strongly inhibit one another while providing little inhibition to other interneurons, in contrast with SST interneurons *(47,* 53).

To mimic the somatic impact of DBS on cortical neurons, we added an external current IDBS to the equation of the voltage variable in every cell of the network. More precisely, considering the periodic nature of DBS-induced somatic currents, I_DBS_ corresponds to a series of square pulses of fixed duration (2 ms) and amplitude (120 pA), followed by a period of silence. These pulses were repeated at a variable frequency, which unless stated otherwise was set to the classical value of 130 Hz. I_DBS_ impact neurons after specific delays, chosen heterogeneous: 0 ms for half of pyramidal cells and PV interneurons, 2 ms for the other half of pyramidal cells and SST interneurons. For optogenetic stimulations, a series of square pulses of duration 3 ms and amplitude 400 pA was applied to half of PV or SST neurons. This fraction was chosen considering the fact that approximately a 70% of targeted neurons express the channelrhodopsin, among which about 70% respond to the light presentation.

Simulations of the network activity were done using a custom code developed in Matlab R2016 (The Mathworks).

### Analysis

*Firing rates:* For each condition, the firing rates of the different neuronal populations were computed as the average rate found over 100 independent simulations (with different intrinsic noises and connectivity patterns) of duration 1.2 s. The averages were computed over the last 1000 ms, to avoid a bias due to transient responses occurring within the first 200 ms. When mimicking PV or SST opto-activation, the average firing rates of PV or SST interneurons were calculated only based on the neurons that were directly activated.

Heatmaps of pyramidal cells’ firing rate modulation were obtained by varying the intensity (from 0 to 120 pA) of *I_Dbs_* received by two populations at the same time, while keeping the default intensity (120 pA) for the third population. When PV and SST receive varying intensities of I_dbs_, only half of the pyramidal cells receives 120 pA DBS currents, while the other half did not receive any *I_DBS_* (Fig. 4C). For the two other cases (Fig. S5), only half of pyramidal cells received varying intensities of I_dbs_, while the other half received 120 pA DBS current pulses. Heatmaps were obtained from the average firing rate of pyramidal cells (found over 20 independent simulations), divided by the average firing rate of pyramidal cells in the parkinsonian condition.

A linear regression of pyramidal cells firing rates as a function of the amount of *I_DBS_*(corresponding to the total amount of current in pA injected over one second, equal to:

stimulation intensity x pulse duration x stimulation frequency) was performed. All three populations received the same inputs, ranging from 40 to 200 pA, with pulse duration lasting from 1 to 3 ms, repeated at 13, 67, 130 and 200 Hz. For each condition, 50 independent simulations were run.

*Pearson correlation coefficients:* To test the capacity of the network to discriminate and respond specifically to various inputs, an additional current was injected to a subset of pyramidal cells (200 randomly chosen cells). We explored the responses of the network to three types of stimuli, presented during 500 ms: constant input, linear ramping input (decreasing or increasing amplitude with time) or noisy inputs (Ornstein-Uhlenbeck processes x_t_ with different mean μ, variance o and time constant τ, according to 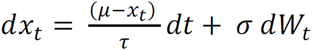 with x_0_ = 0). These either deterministic or stochastic inputs (Table S2) were chosen such as to mimic some activity patterns observed in pyramidal neurons in the motor cortex (55).

For each condition (control, Parkinson, Parkinson+DBS at 130Hz, Parkinson + opto-activation of SST or PV interneurons at 67Hz), we used two different methods for quantifying the capacities of extracting information from the network activity patterns.

We first measured for each trial the Pearson correlation coefficient ρ between the moving spike count of all pyramidal cells across time (response: R) for ramping and stochastic stimuli (stimulus: S), defined according to the following formula: 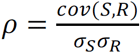 with *cov*, corresponding to the covariance and *σ,* to the standard deviation. The moving spike count was calculated based on the sum of the number of spikes across all pyramidal cells, with a moving time interval of 10 ms. The first and last 10 ms of stimulus presentation were discarded to avoid boundary effects. For each stimulus, an average correlation coefficient was obtained by averaging the Pearson correlation index across 100 independent trials (with different intrinsic noises, but identical connectivity matrices, to explore specifically the variability of the responses of a given network to the same input). Example trace of this moving spike count for 5 different trials (obtained from the control condition on stimulus n°4) and the resulting distribution of correlation coefficient over all trials are shown in Fig. 4F.

*Classification of stimuli:* We also estimated the efficiency with which downstream neurons might discriminate the network responses to various stimuli, beyond the sole knowledge of the mean firing rate. We used supervised-learning algorithms to classify the output of the above-described network. Our approach consisted of the following steps (Fig. 4F). We first selected the responses of the network to 20 stimuli (the four constant stimuli, the first eight ramping stimuli and first eight stochastic stimuli; Table S2). We considered as our network responses a time-binned matrix M in which each row corresponding to a given pyramidal cell would contain the number of spikes emitted for every 10 ms. The first 200 rows corresponded to the responses of the pyramidal cells directly activated by the stimulus. In order to reproduce the highly convergent cortical motor inputs received by striatal neurons, we densified the response of pyramidal cells by contracting these 800×50 M matrices into a 100×50 S matrix, defined as: S=W.M, where the weight matrix W is a random matrix, identical for all stimuli presentations, with each element generated from the uniform distribution on the interval [0,1] (Fig. 4F). These S matrices were then used as the input for the supervised-learning algorithms. We used four classical algorithms: nearest centroid classifier, multinomial logistic regression, linear discriminant analysis and support vector machines (Table S3). We refer to Bishop (56) and Hastie et al. *(57)* for conceptual details on these algorithms. For each condition, the dataset consisted of 100 repetitions for each of the 20 stimuli (with the same network parameters and convergence matrix W, but independent realizations of the intrinsic noise £,). The four classifiers were trained to discriminate the population response given the stimulus on 80% of the data sample, using stratified k-fold cross validation (k=5). We defined the accuracy of the classifiers as the percentage of correct predictions of the stimulus identity from the network response. The training and testing of the classifiers were run using *scikit-learn* and *keras* packages in Python 3.5 (Python Software Foundation, www.python.org).

*Statistics:* Normally distributed data are displayed as mean ±SD, and differences between conditions (control, Parkinson, Parkinson+DBS, opto-activation of SST or PV interneurons at 67 Hz) were assessed using one-way ANOVA for each feature and Tukey-Kramer *post-hoc* test. Since some distributions of correlation coefficients were not normal, all data is summarized using the median and presented in details as boxplots (using the same convention as described above) and differences between conditions were assessed using the Kruskal-Wallis test (with Tukey-Kramer *post-hoc* test) on every stimulus.

---

**Supplementary Figure 1.**
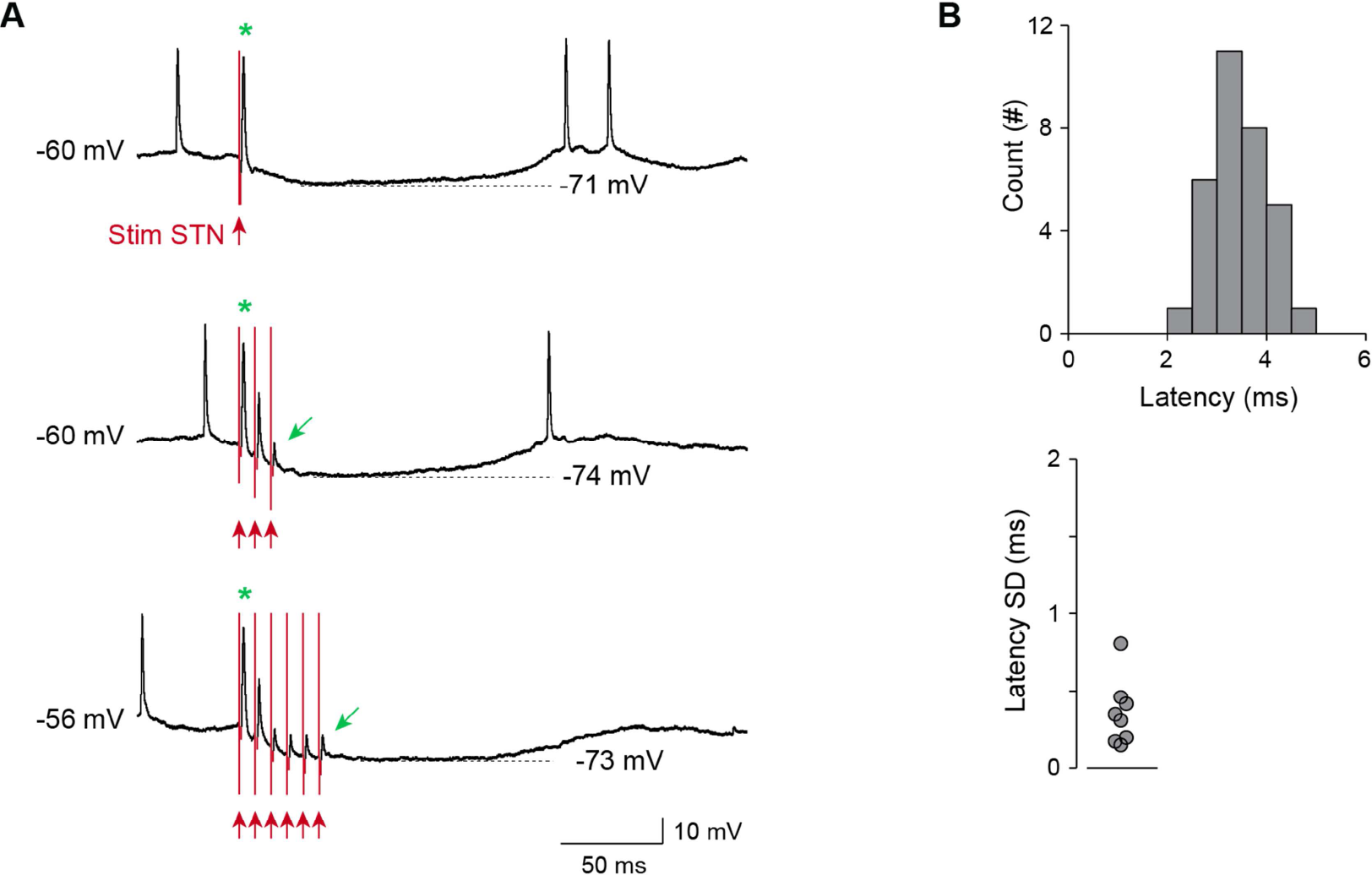
STN single stimulation in M1 pyramidal cells. (A) Example of an intracellularly recorded M1 pyramidal cell antidromically activated by STN stimulation (red arrow: STN stimulation; green star: antidromically evoked spike). These cells (n=4) also displayed a pause in firing after the antidromic spike, associated with an hyperpolarization of the membrane potential (upper panel). Upon repetitive STN stimulations at 130 Hz (middle and lower panels), the antidromic spike amplitude is dramatically reduced and the amplitude of the hyperpolarization is increased. (B) Distribution of the latency of the response evoked by STN single stimulation in M1 pyramidal cells (n=8 neurons, 4 measures per neuron) and the SD of the response latency for each cell.

**Supplementary Figure 2.**
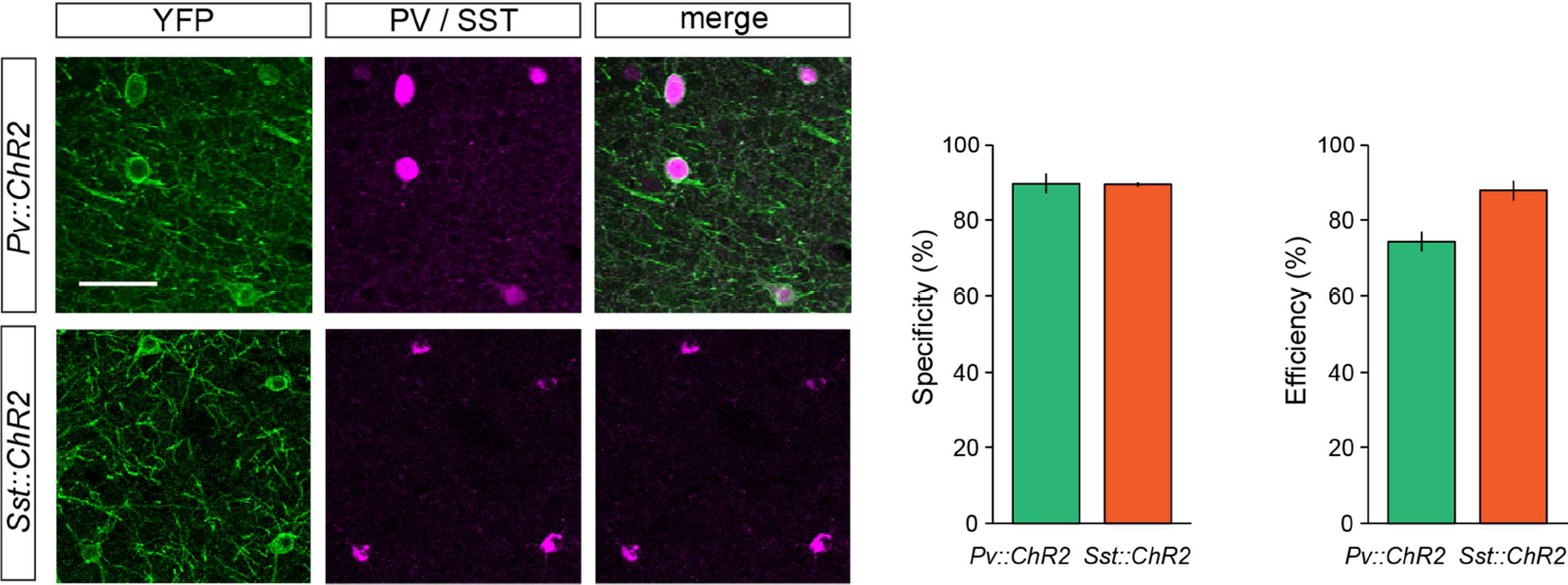
Specific expression of ChR2 expression in cortical PV and SST interneurons. Representative confocal pictures of M1 cortex in *Pv::ChR2* (top) and *Sst::ChR2* (bottom) mice, with double immunostaining for YFP (ChR2 expression reporter, green), and either PV (for *Pv::ChR2* mice) or SST (for *Sst::ChR2)* (pink) (right panels). Quantification of the specificity (number of double labeled cells / number of YFP-positive cells) and efficiency (number of double-labeled cells / number of PV-or SST-positive cells) of ChR2 expression in *Pv::ChR2* and *Sst::ChR2* mice (mean±SEM of 4 hemispheres from 2 mice in each mouse strain) (left panels). Scale bar: 50 pm.

**Supplementary Figure 3.**
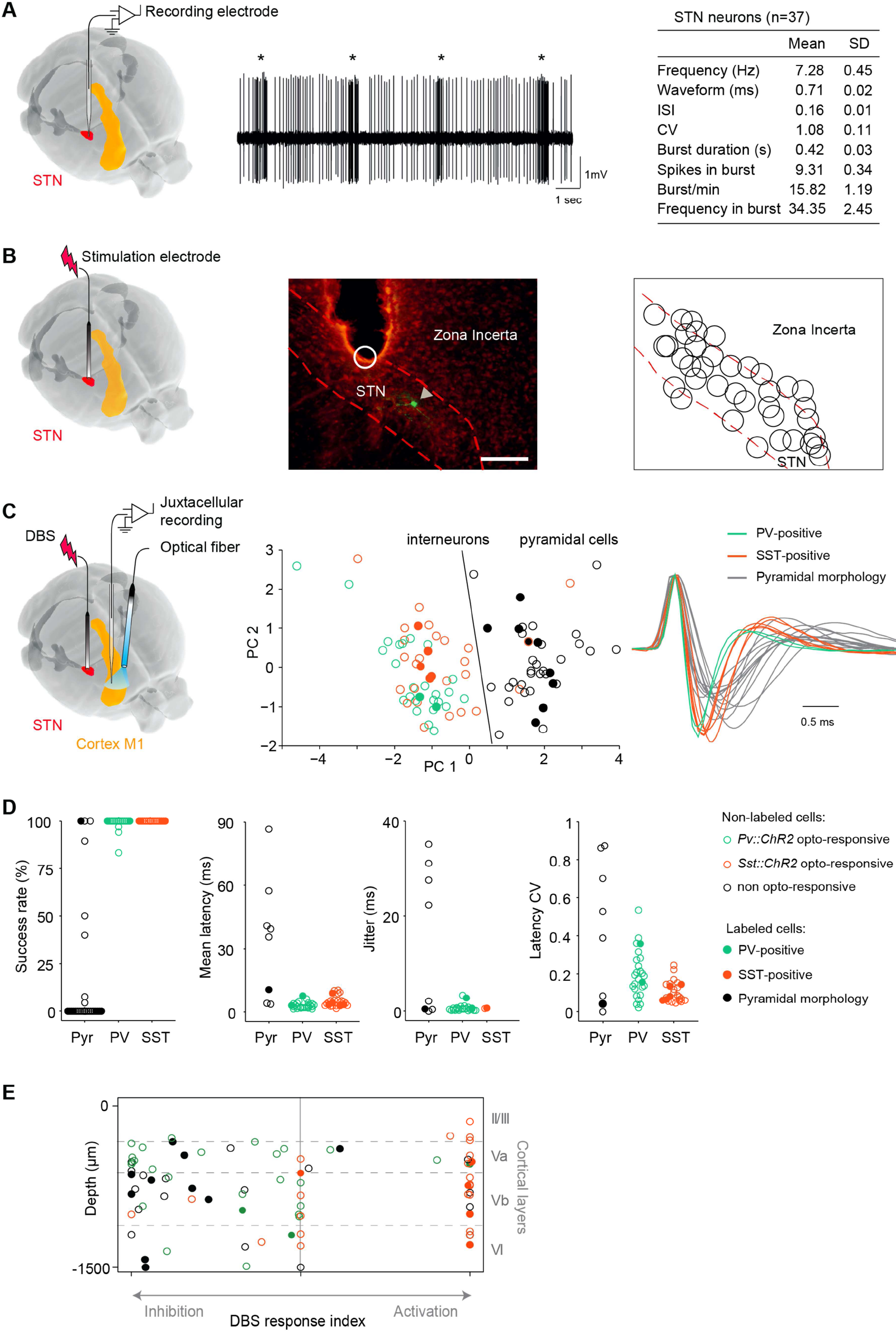
Methodology for STN targeting and identification of M1 neurons. (A) Electrophysiological identification of STN position. Left, a microelectrode is lowered in a stereotaxically defined region until encountering a neuron with typical STN firing patterns. Middle, extracellular recording of an STN neuron (stars indicate bursts as detected by the Poisson surprise method). Right, table represents the electrophysiological characteristics of STN neurons (n=37). (B) The DBS stimulation electrode is lowered in the same coordinates as the identified STN neurons. Middle, photomicrograph shows a juxtacellularly labeled STN neuron (white arrow) in close proximity to the tip of a stimulating electrode (white circle). Scale bar=250μm. Right, schematic representation of the location of all electrically induced lesions of the STN. All data obtained with lesions outside of the STN were not included in the study. (C) With the DBS electrode in place, an optical fiber is placed on top of the M1 cortex and a microelectrode is lowered close to the optical fiber until encountering a photo-responsive neuron. Middle, principal component analysis of waveform characteristics and opto-response properties reveal cutoff boundaries for discriminating pyramidal neurons from interneurons. Five opto-responsive neurons with non-matching morphology and/or waveform were excluded. Right, waveforms of all labeled neurons (SST in orange, PV in green and pyramidal neuron in grey. Dashed waveforms correspond to excluded neurons). (D) Success rate, mean latency, jitter and latency CV of the response to opto-activation in all neurons (pyramidal neurons in black, PV in green and SST in red; immunohistochemically or morphologically identified neurons are represented as filled circles). (E) Cortical depth of all recorded neurons (based on position of labeled neurons or on manipulator depth) plotted according to their response to STN-DBS.

**Supplementary Figure 4.**
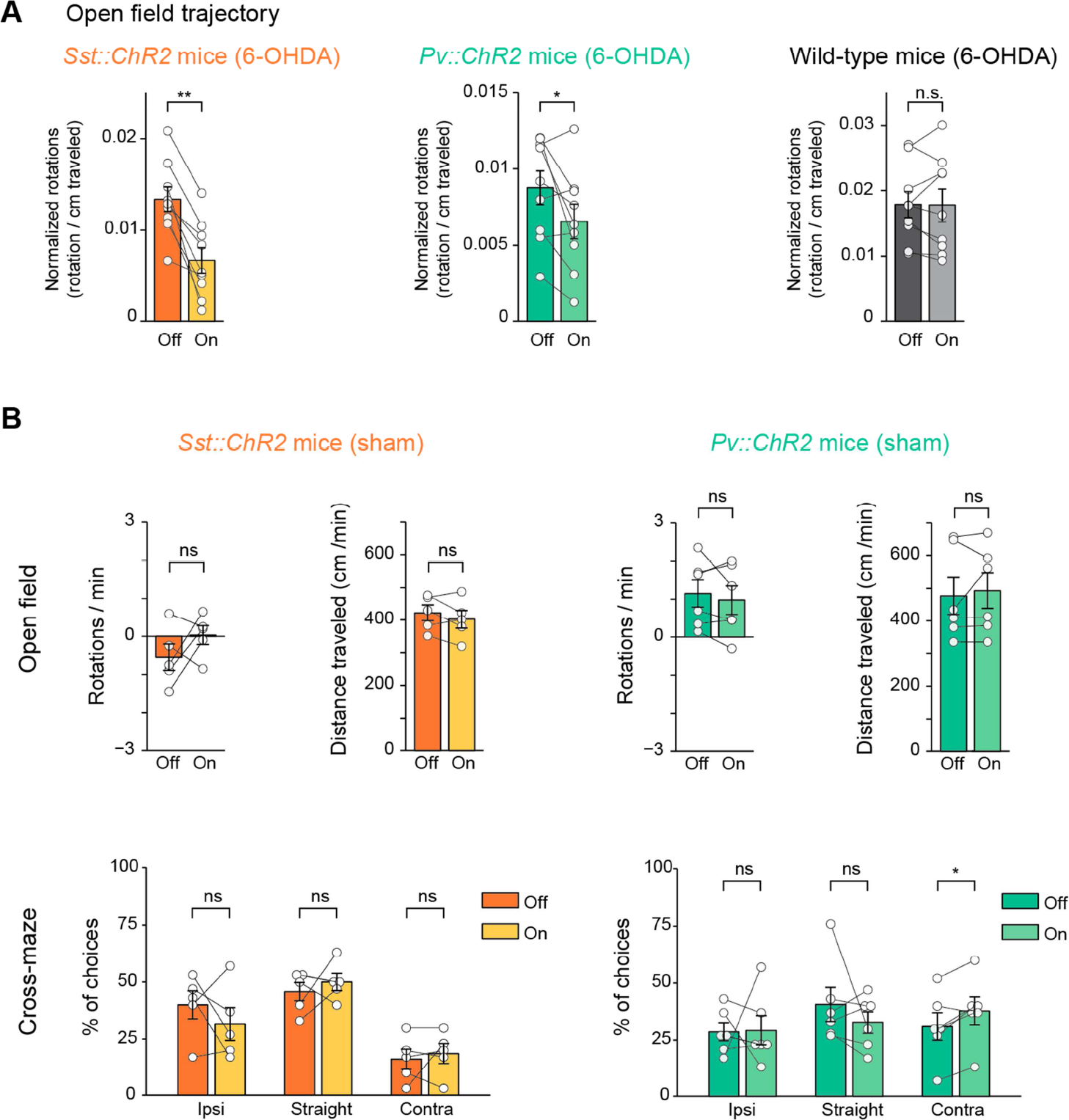
Additional results related to the Figure 3. (A) In the open field, the effect of light on rotational behavior was still present when rotations were normalized by the distance travelled in each light-off or light-on epoch, inducing a decrease in 6-OHDA *Sst::ChR2* (n=9, p<0.001, paired t-test) and 6-OHDA *Pv::ChR2* (p=0.042) mice, but not in 6-OHDA WT mice (p=0.92). (B) In the open field task, opto-activation of PV or SST interneurons in sham mice does not affect rotational behavioral and locomotor activity *(Pv::ChR2* : n=6, p=0.43 and p=0.55; *Sst::ChR2: n=5, p=0.25* and p=0.27). In the cross-maze task, opto-activation of PV or SST interneurons in sham mice does not lead to preferential turn *(Pv::ChR2* : n=6, ipsilateral turns p=0.94, straight p=0.39 and contralateral turns p=0.027; *Sst::ChR2: n=5,* ipsilateral turnsp=0.37, straightp=0.53 and contralateral turnsp=0.61).

**Supplementary Figure 5.**
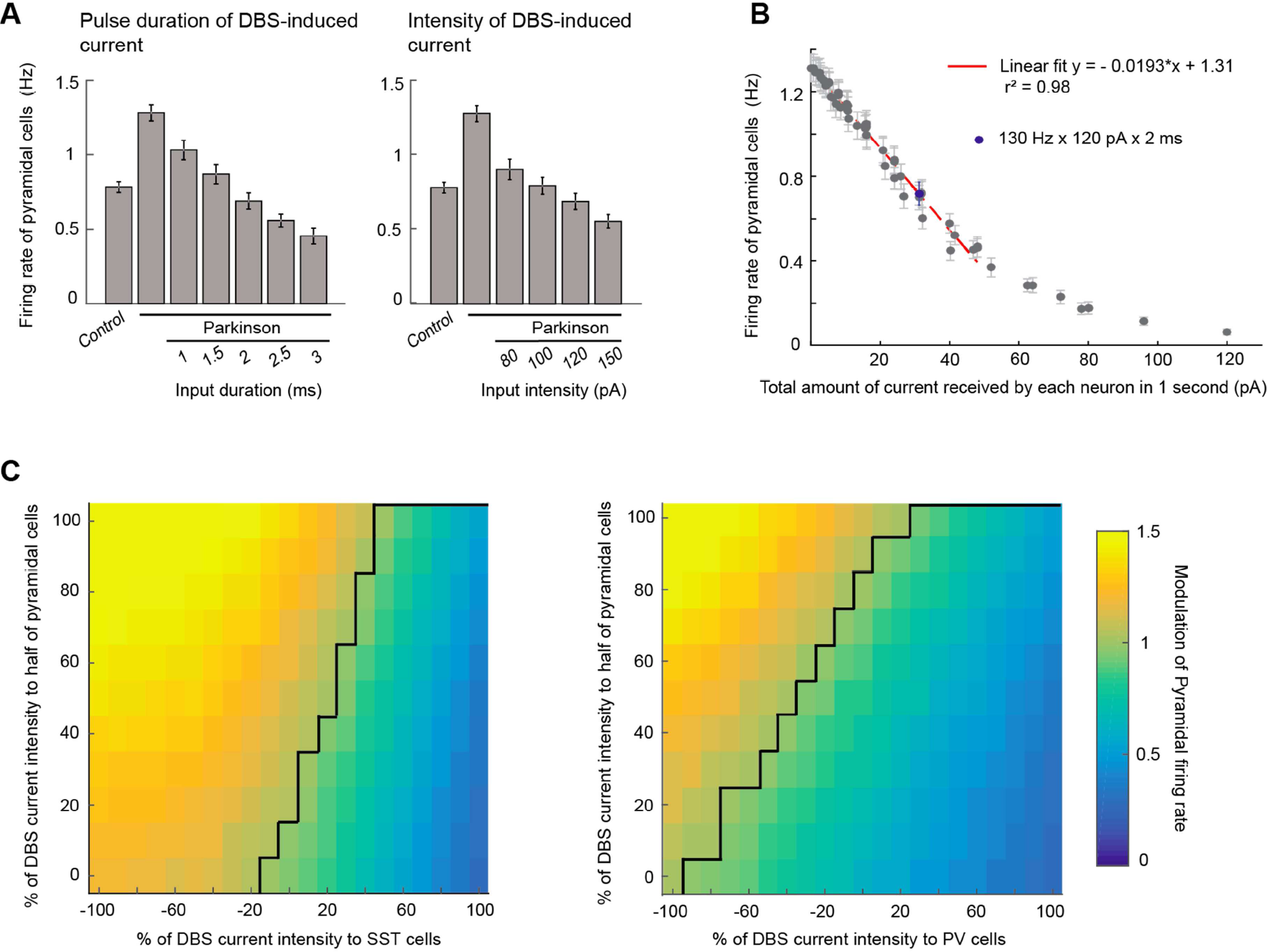
Estimating the robustness to changes in DBS parameters. (A) Average firing rates (±SD) of pyramidal cells when varying two parameters of DBS-induced currents for all three populations (pulse duration and intensity). All conditions were significantly different from the parkinsonian condition (p<0.001). (B) Evolution of pyramidal cell average firing rate (±SD) as a function of the amount of DBS-induced current injected in the network (total current received by each neuron over one second). A linear fit (red dashed line) was performed, without considering the last eight points corresponding to high intensity currents. (C) Average firing rate of pyramidal cells relative to the parkinsonian condition as a function of the percentage of DBS current intensity received by half of pyramidal cells, all SST or PV interneurons. The maximal intensity (100 %) corresponds to 120 pA. The other half of pyramidal cells receives a 120 pA DBS-induced current in all cases. The isocline corresponding to a ratio of 1 is indicated with a black line.

**Supplementary Figure 6.**
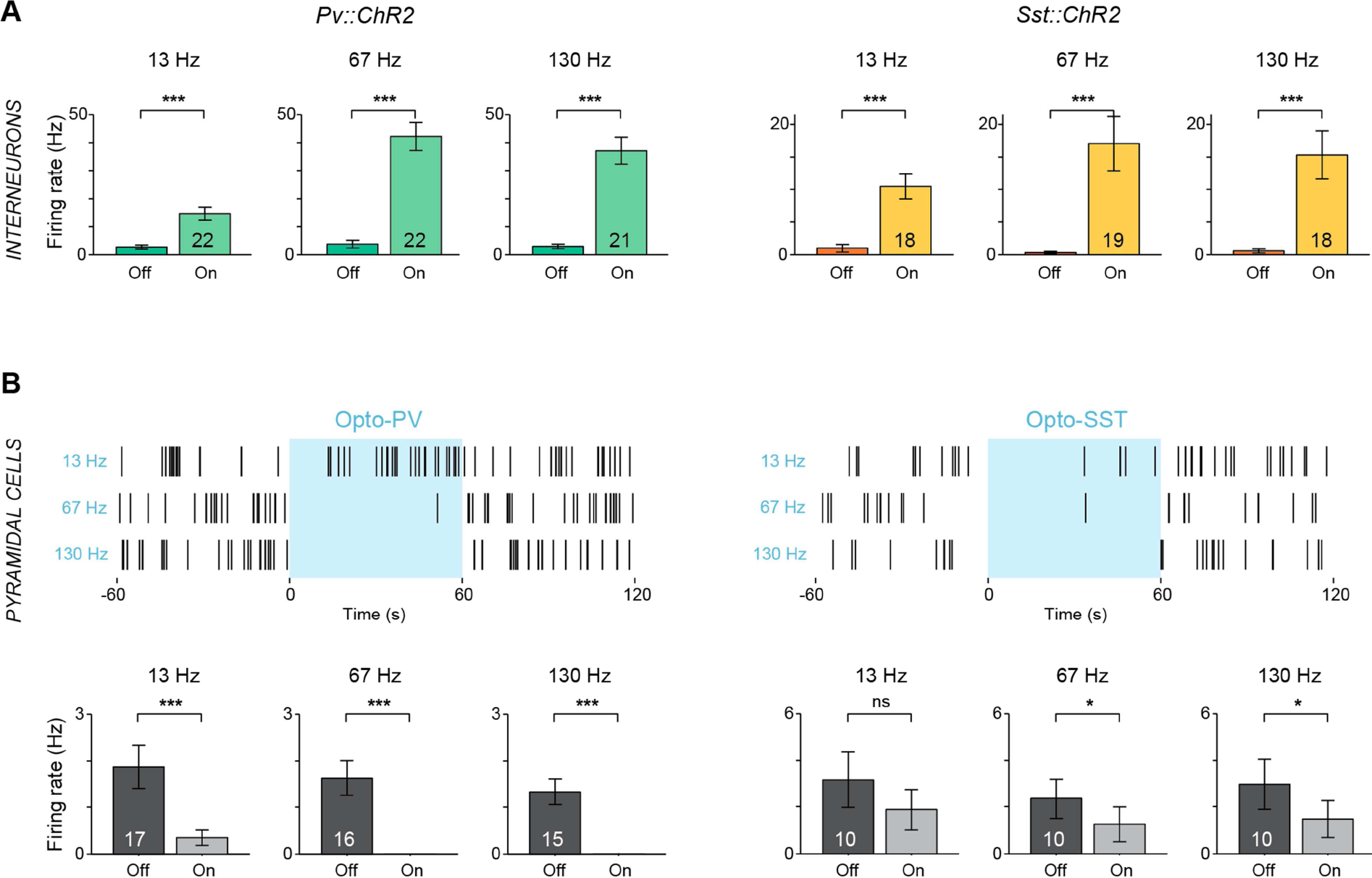
*In vivo* **effect of opto-activation of PV or SST interneurons.** (A) Opto-activation of M1 PV (left) and SST (right) interneurons (3 ms pulses, 10 mW at 13, 67, or 130 Hz for 1 min). Mean±SEM, ***: p<0.001 (Wilcoxon’s signed rank test). (B) Upper panels: rasterplot of the activity of two pyramidal cells recorded during opto-activation of PV (left) or SST (right) interneurons at 13, 67 or 130 Hz. Bottom panels: pyramidal cell activity is decreased upon activation of PV (left) or SST (right). Mean±SEM, *: p<0.05, ***: p<0.001 (Wilcoxon’s signed rank test).

**Supplementary Figure 7.**
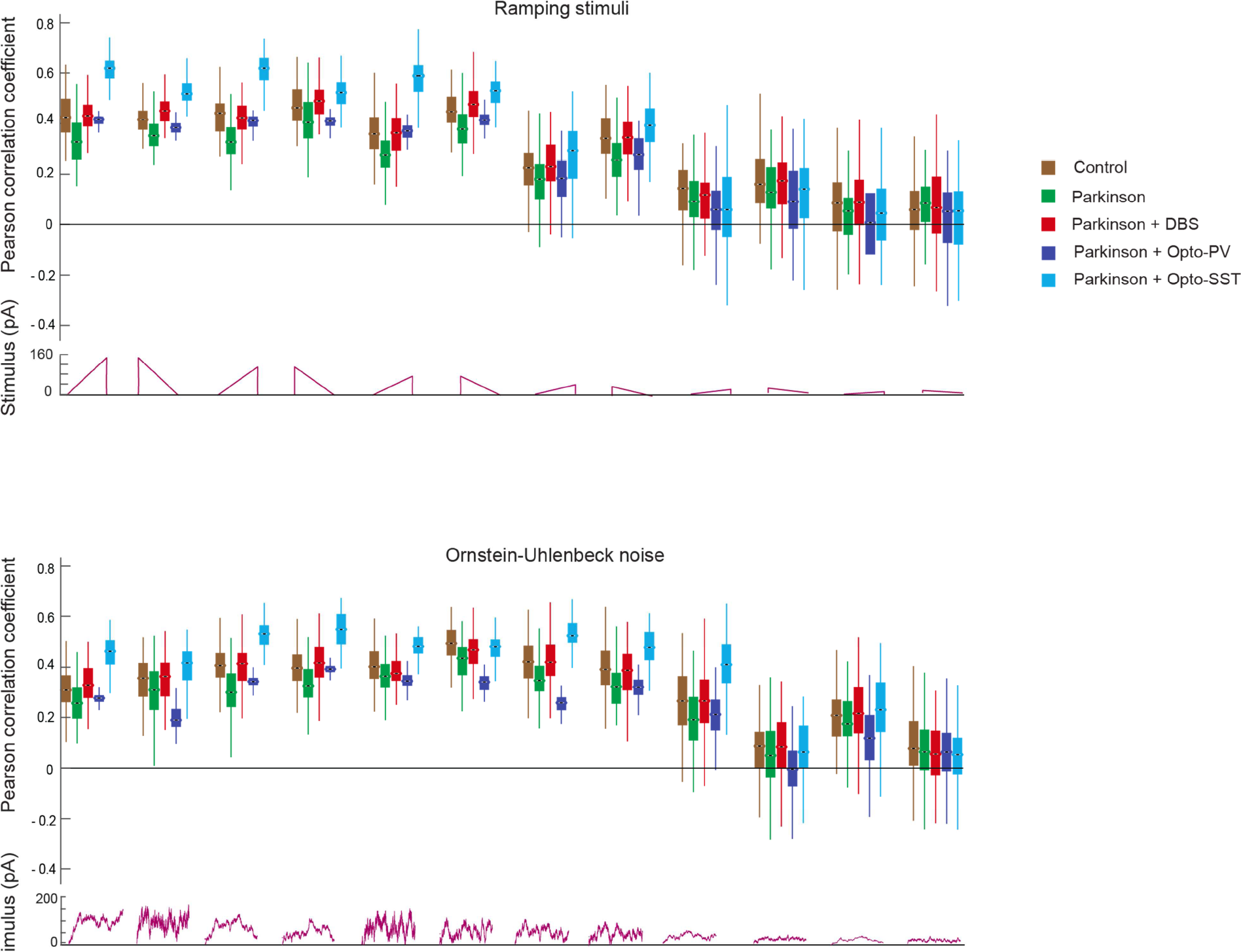
Correlations between the network responses and time-varying stimuli. Pearson correlation coefficients between an input (displayed below) and the moving spike count over all pyramidal cells (with a 10 ms interval), for twelve ramping stimuli *(top)* or Ornstein-Uhlenbeck processes *(bottom).* Statistical differences are indicated in Fig. 4G.

**Supplementary Table 1.**
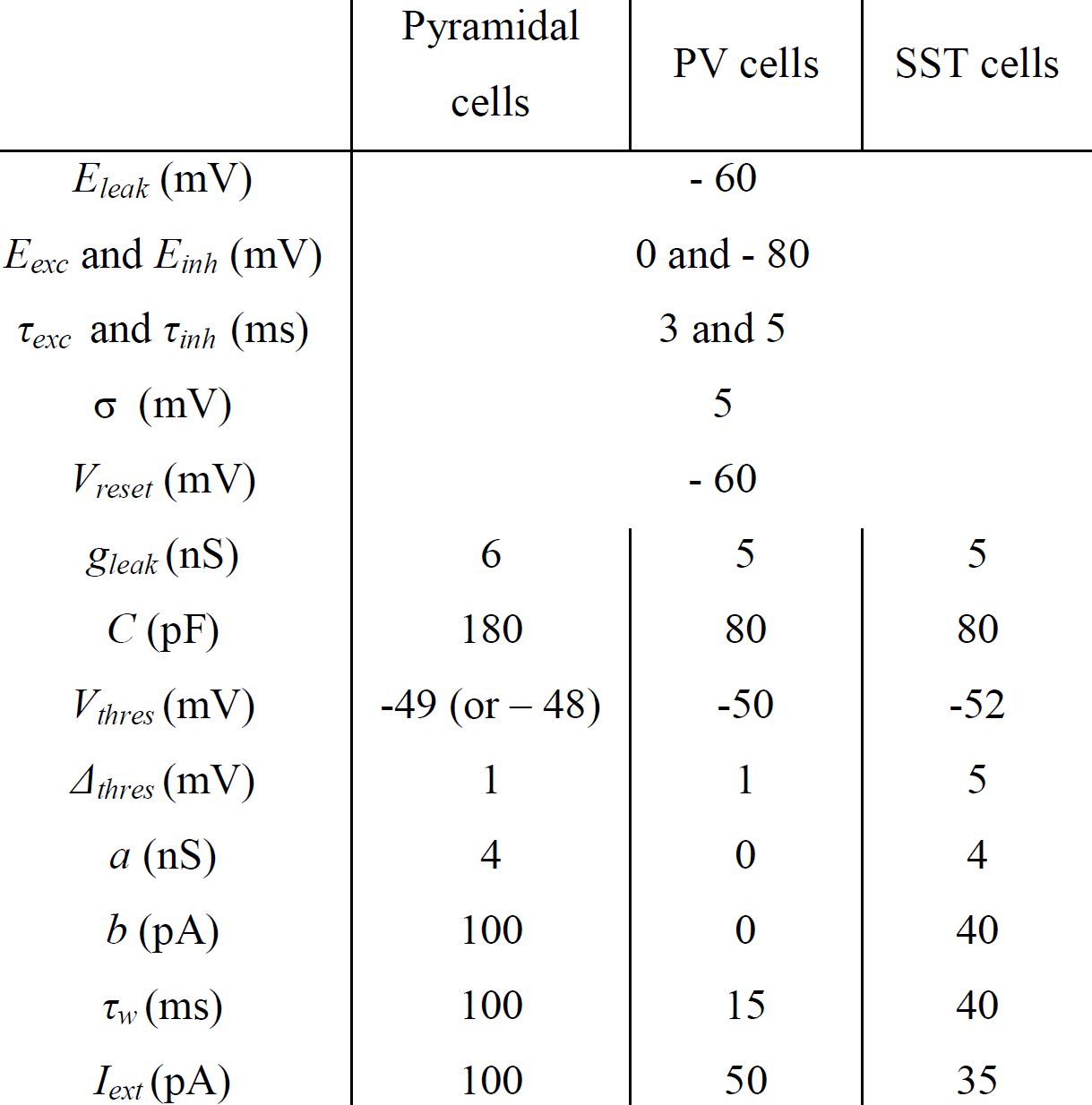
Intrinsic electrophysiological parameters for each neuronal subtype.

**Supplementary Table 2.**
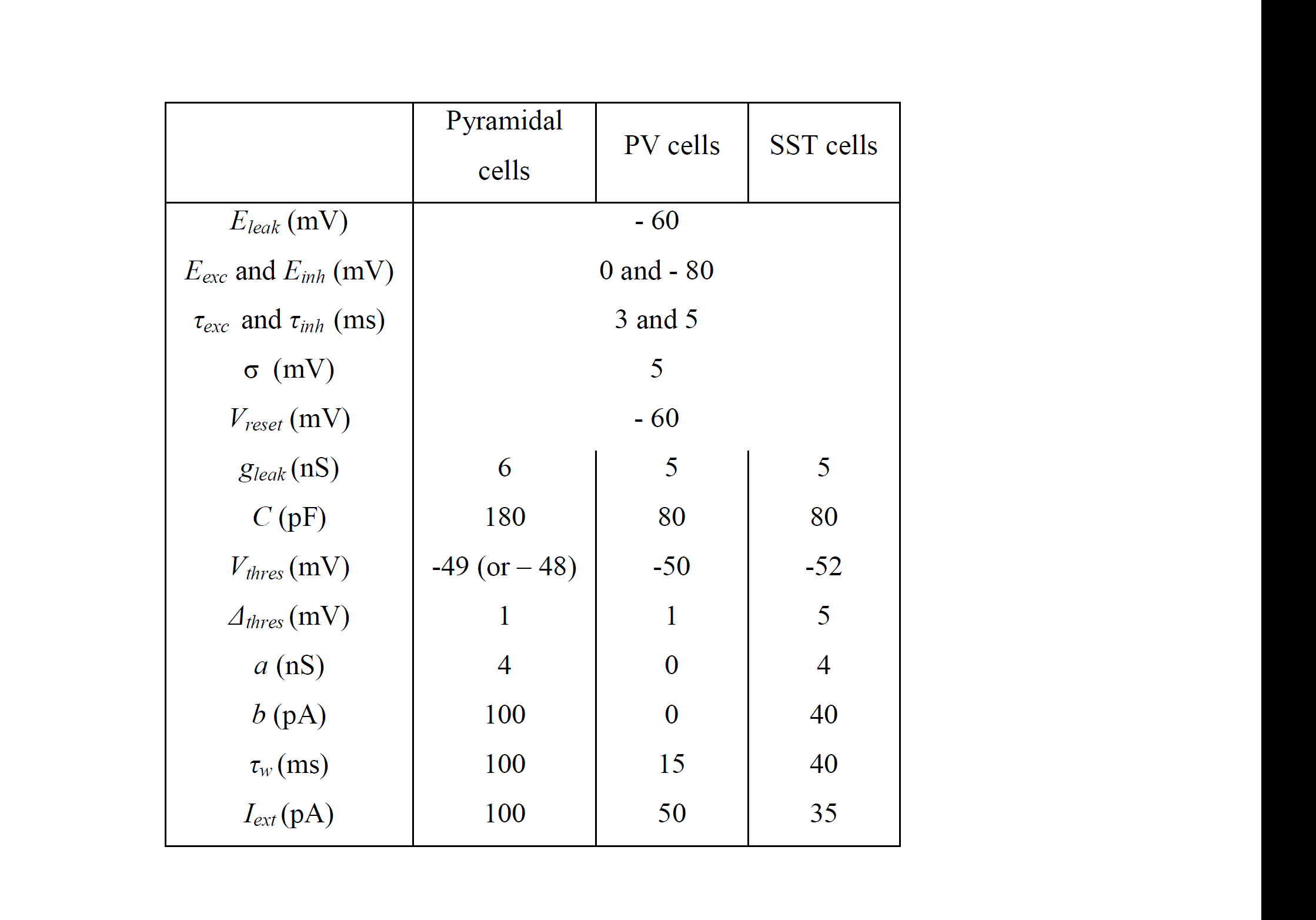
Choice of stimuli injected to a subset of pyramidal cells.

**Supplementary Table 3.** Details of the functions and parameters used for each supervised-learning algorithm.

| Algorithms | Functions and Parameters |
| --- | --- |
| Nearest centroid classifier | Use of the Euclidean distance<br>No threshold for shrinking centroids |
| Multinomial logistic regression | Use of the cross-entropy loss for the decision function<br>Using the Newton-cg solver (with L2 regularization)<br>Penalty of the error term: C=1 (default parameter)<br>Tolerance: 0.00001 (default parameter)<br>Maximal number of iterations: 300 |
| Linear discriminant analysis | Using singular value decomposition<br>No priors given, no given knowledge of the number of components for dimensionality reduction<br>Tolerance: 0.0001 (default parameter) |
| Support vector machines | Use of a “one-against-one” approach for the decision function<br>Using a linear kernel, with a shrinking method<br>Penalty of the error term: C=1 (default parameter)<br>Tolerance: 0.0001 (default parameter)<br>No limit on the number of iterations |

